# Quantitative comparison of fungal genome assembly strategies using short and long-reads from simulated and empirical sequencing data

**DOI:** 10.64898/2026.06.03.729889

**Authors:** Gabriel A. A. Silva, Temitope R. Folorunso, Lori G. Eckhardt, Janna R. Willoughby

## Abstract

High-quality fungal reference genomes are essential for comparative, functional, and evolutionary studies, yet fungal genome features such as repeats, structural rearrangements, accessory chromosomes, and intron-rich genes can complicate genome assembly and the selection of cost-effective sequencing strategies. Here, we benchmark fungal genome assembly performance using simulated and empirical short- and long-read datasets to evaluate how sequencing depth, assembler choice, and genome characteristics influence contiguity, completeness, accuracy, and computational requirements. Using simulated reads from complete fungal genomes spanning diverse sizes and compositions, we evaluated short-read, long-read, hybrid, and polished long-read assemblies across sequencing depths from 10X to 100X. Key trends were validated using empirical sequencing data from 10 fungal isolates assembled with multiple strategies, including different Flye assembler parameter sensitivity and short-read polishing. Across datasets, long reads produced the largest improvements in contiguity, with most gains achieved at ∼20-40X coverage and diminishing returns beyond moderate depth. Short-read polishing substantially improved base-level accuracy at relatively low cost, with ∼10-20X coverage often sufficient to approach maximal error reduction. Hybrid assemblers showed strong algorithmic variability, with trade-offs between contiguity, error rates, and computational demand. Genome architecture also influenced outcomes, as larger and more feature-dense genomes benefited more from long-read data while GC content had limited impact. Overall, our results suggest that moderate long-read coverage (∼30-40X) combined with modest short-read polishing (∼10-20X), particularly using Flye plus Polypolish, provides a strong balance of contiguity, completeness, accuracy, and resource efficiency for generating high-quality fungal genome assemblies.

**Impact statement:** Fungal genome sequencing is expanding rapidly across ecology, plant pathology, biotechnology, and clinical and veterinary microbiology, yet experimental design decisions regarding sequencing depth, assembler selection, and hybrid workflows are still largely guided by bacterial benchmarking studies or limited single-species comparisons. Because fungal genomes vary widely in size, repeat content, and gene architecture, these assumptions can lead to inefficient sequencing strategies, increased computational costs, and suboptimal assemblies. Here, we develop a reproducible assembly benchmarking framework that combines large-scale simulations from 66 complete fungal genomes spanning plant, animal, and human-associated taxa with newly generated short- and long-read sequencing data from 10 field-collected isolates. This approach enables evaluation of assembler performance across diverse genome architectures and tests whether patterns identified in simulations translate to real biological datasets. Across both simulated and empirical datasets, we show that reliable fungal genome reconstruction can be achieved without excessive sequencing depth by identifying consistent performance thresholds. Assembly contiguity and completeness stabilize at moderate long-read coverage, after which improvements depend more strongly on assembler choice and genome structure than on additional data volume. Hybrid workflows show trade-offs in accuracy, contiguity, and computational demand, whereas targeted short-read polishing provides an efficient strategy for improving base-level accuracy. These findings offer practical guidance for fungal genome assembly and support more robust downstream genomic analyses across non-model microbial systems, including ecologically, agriculturally, and clinically important fungi.

**Data summary:** The reference fungal genomes used for simulation are available in the NCBI Assembly database under the accession numbers listed in Supplementary Table S1. Empirical raw sequencing data generated for this study are deposited in the NCBI Sequence Read Archive (SRA) under accession PRJNA1474061.

All scripts used for read simulation, assembly, polishing, benchmarking, and statistical analyses are available in the project GitHub repository (github.com/bielasilva/fungi_assembly_benchmarking). Software versions, parameters, and workflow configurations are provided within the repository and detailed in the Methods.

## Introduction

Fungi are one of the most diverse and socioeconomically important groups of organisms, influencing biotechnology, agriculture, medicine, and environmental science [1]. Their impacts span the production of pharmaceuticals such as antibiotics, statins, and immunosuppressants [2,3], industrial enzymes [4] and biocontrol agents [5], as well as negative impacts as pathogens that cause major losses in agriculture [6], timber industry [7] and livestock [8,9], and pose health concerns for humans [10,11], domestic animals [12] and natural forest ecosystems [13]. Given this ecological breadth and economic impact, elucidating how they respond to environmental stressors, interact with other species, and evolve is fundamental to their sustainable use and management. High-quality reference genomes provide the foundation for these insights, enabling comparative, functional, and evolutionary analyses across this extraordinarily diverse kingdom, and support clinical applications such as pathogen identification, antifungal resistance tracking, and outbreak surveillance.

Despite their importance, high-quality reference genomes remain scarce for many fungal taxa, particularly among ecologically important plant pathogens, endophytes, and soil-associated lineages where assemblies are often fragmented or unavailable [14,15]. Although typically smaller than those of plants and animals, fungal genomes can greatly vary in size, organization, and gene content [16]. Fungal genomes range from compact, haploid genomes of less than 10 Mb to large, repetitive, and polyploid genomes exceeding 1 Gb [15]. Examples at the small end include the human-associated pathogens *Malassezia restricta* [17] and *Pneumocystis jirovecii* [18], while large-genome species include the maize rust *Puccinia polysora* [19] and the cicada-infecting *Massospora cicadina* [20]. In addition, many species possess accessory or conditionally dispensable chromosomes that contribute to host specificity, virulence, or environmental adaptation [21,22]. Frequent structural rearrangements [23,24], abundance of transposable elements [25], horizontal gene transfers [26] and complex gene clusters [27] create computational challenges to generating high quality reference assemblies in these fungal species.

Because of the genomic complexity, the accuracy and completeness of fungal genome assemblies depend on the sequencing technology and assembly strategy. Traditionally, the choice of sequencing technology determines the assembly outcome and downstream analytical options. Illumina short-read sequencing has long been the gold standard for base-level accuracy (<0.1% error rates), but at the cost of limited resolution potential in repetitive regions, leading to fragmented assemblies [28]. In contrast, Oxford Nanopore Technologies long-reads can exceed 100 kb, facilitating the resolution of complex repetitive elements and structural variants. Earlier nanopore chemistries had error rates exceeding 5-10%,which historically limited their use for high-accuracy applications [29], but successive improvements in pore chemistry, basecalling models, and read length have substantially reduced raw-read error rates and yielded high consensus accuracy on bacterial genomes, comparable to or exceeding short-read assemblies [30–32].

Given the historical limitations of single-platform assemblies, hybrid approaches have emerged as an alternative to the challenges of single technology assemblies. These methods combine the contiguity advantages of long reads with the base-level accuracy of short reads [33], and post-assembly polishing tools, such as Pilon [34] and Polypolish [35], using short reads can significantly improve the base-level accuracy of long-read assemblies [36]. However, the optimal combination of assemblers, polishing tools, and sequencing depths for fungal genomes remains poorly explored. Previous benchmarking studies have primarily focused on bacteria [36–39], with the few exceptions targeting single-taxon model eukaryotic systems (e.g., *Giardia* [40], *Saccharomyces* [41], *Drosophila* and human reads [42]) and limited subsets of assemblers and sequencing depths, leaving the distinctive challenges of fungal genome architecture largely unaddressed. This warrants a systematic, large-scale evaluation to guide cost-effective experimental design and accurate genome reconstruction.

This study addresses these critical knowledge gaps through a systematic comparison of fungal genome assembly approaches. First, we considered how sequencing strategy and coverage depth influence assembly size, contiguity, completeness, accuracy, and computational cost in fungal genomes across four distinct strategies sequencing and assembly strategies: short-reads-only, long-reads-only, hybrid, and long-reads-polished, evaluating across multiple sequencing depths (10X to 100X coverage) to determine optimal resource allocation strategies. Next, we considered how intrinsic genome characteristics of 66 fungal pathogens, including genome size, number of genes and coding elements modulated assembler performance. Finally, we considered which combinations of sequencing technology, depth, and assembly workflow supported the most efficient trade-offs between quality and computational resources to provide evidence-based recommendations for fungal genome assembly workflows. This was designed to enable researchers to make informed decisions about sequencing strategies and computational pipelines tailored to their research and resource availability.

## Methods

### Overview

We evaluated fungal genome assembly performance using a two-part framework that combined simulations with empirical validation. First, we measured assembly behavior using simulated short- and long-read sequencing data from complete fungal reference genomes and generated assemblies across multiple sequencing depths with representative short-reads-only, long-reads-only, hybrid, and long-read-plus-polishing strategies. These assemblies were compared using standardized metrics of contiguity, completeness, base-level accuracy, assembly size, and computational resource requirements, and genome characteristics were incorporated to examine how intrinsic features influenced performance. Second, to evaluate how these patterns translated to empirical datasets, we sequenced 10 distinct fungal isolates with both short- and long-read technologies and applied a focused subset of assembly workflows to assess depth-dependent trends. These isolates were treated as independent empirical genomes for benchmarking purposes, and species-level taxonomic assignments were not required for the assembly-performance comparisons and were not used as a factor in the analyses.

### Simulated genome data

We simulated short-reads and long-reads sequencing data from 66 complete fungal genomes, representing all available assemblies on NCBI RefSeq as of July 29, 2025 (Supplemental Table S1). These reference genomes represented a range of taxa and genomic complexities, so we summarized key characteristics of the reference assemblies by quantifying GC content, total gene count, number of contigs and total assembly size. Assembly size, GC content, and number of contigs were quantified by custom scripts and gene counts were predicted using GeneMark-ES [43]. Reads for each assembly were simulated using NanoSim v3.2.3 [44] for Nanopore and InSilicoSeq v2.0.1 [45] for Illumina (2x250 bp) at 100X coverage. Subsequently, seqtk v1.5 [46] was used to subsample the reads to 10X, 15X, 20X, 25X, 30X, 35X, 40X, 50X, 60X and 75X coverage.

Using the simulated reads, including down sampled datasets, we considered four complementary assembly strategies and compared performance among these approaches. We considered short-reads-only assemblers SPAdes v4.2.0 [47] and ABySS v2.3.10 [48] as well as long-reads-only assemblers Flye v2.9.6 [49], Canu v2.3 [50], Hifiasm v0.25.0 [51], and Raven v1.8.3 [52]. Following these, we considered hybrid assemblers ABySS Hybrid v2.3.10 [48], MaSuRCA v4.1.4 [53], and hybridSPAdes v4.2.0 [47]. Finally, we used the top-performing long-reads-only assembler to be short-read polished by Pilon [34], Polypolish [35], or Racon [54]. Within this set, Racon was run with both bwa-mem2 v2.3 [55] or Minimap2 v2.30 [56] as mappers to assess the impact of the mapper on polishing. Each assembler or polisher was run closest to the recommended parameters and standardized computational resources (*-threads 10* or equivalent) to ensure comparability, and full scripts can be found on GitHub. Canu was also run with the *genomeSize* parameter set to the known genome size, as it is a required parameter.

### Empirical genome data

For the empirical data assessment, 10 distinct fungal isolates were obtained from symptomatic pine needles collected from diseased trees in pine forests in the southeastern United States. These isolates were selected to provide independent empirical genome assemblies for workflow validation, as species-level taxonomic assignments were not used in the benchmarking analyses, and we therefore refer to them here as fungal isolates rather than as species. Sampling, isolation, and culturing followed established protocols described in Folorunso et al. [57]. In short, infected needles were surface-sterilized, sectioned, and plated on Malt Extract Agar medium to obtain pure fungal cultures. The isolates were cultured in liquid Yeast Extract Peptone Dextrose medium, the mycelia were collected, and the DNA was extracted using a modified cetyltrimethylammonium bromide (CTAB) protocol optimized for fungal tissues rich in secondary metabolites and polysaccharides. The full step-by-step protocol is available in Folorunso et al. [57] and archived on Protocols.io [58].

Following the simulated data approach, we generated short- and long-read sequencing data for each of the 10 fungal isolates. For short-reads, sequencing was carried as 150bp PE on Element Biosciences AVITI24 by the University of Minnesota Genomics Center. Raw short-reads were processed using Fastp v0.24.1 [59] *(--correction --detect_adapter_for_pe --trim_front1 20 --trim_front2 20 --trim_tail1 2 --trim_tail2 2*). For long-reads, we used the Oxford Nanopore Technologies MinION Mk1C or Mk1D, the Native Ligation Barcoding Kit 24 (SQK-NBD114.24) on flow cells R10.4.1 (FLO-MIN114). We basecalled with Dorado v0.9.1 (super-accuracy model v5.0.0) and filtered long reads with Chopper v0.11.0 [60]. No fixed number of bases were removed from the ends of reads, and filtering was limited to removing reads shorter than 1,000 bp (*--minlength 1000*). Based on simulated data output, we assembled using ABySS, SPAdes (short-reads only and hybrid), Flye (with *--min-overlap* set to 1000, 1500, 2000, 2500 or 3000) and Flye + Polypolish.

### Assembly comparisons

For simulated datasets, assembly performance was evaluated using reference-based metrics. Total assembly length, N50, number of contigs, and base-level accuracy were quantified with QUAST v5.3.0 [61] and assembly completeness was assessed with BUSCO v6.0.0 [62] using the fungi_odb10 lineage dataset. Relative assembly size was calculated as the difference between the assembled genome length and the corresponding reference genome used to generate simulated reads, enabling standardized comparisons across assemblers and sequencing depths. Similarly, contiguity differences were assessed by comparing the number of contigs produced by each assembly to those of the reference genome. Because the true genome sequence was known, BUSCO completeness, genome fraction recovered, length error, and base-level error rates were interpreted as direct measures of assembly accuracy relative to the reference.

For empirical datasets, where a ground-truth genome was unavailable, assemblies were compared using reference-free assembly metrics, k-mer-based genome size estimates, and BUSCO gene completeness. Genome size was estimated independently from both short-read and long-read datasets using KMC v3.2.4 [63], followed by GenomeScope2 v2.0.1 [64]. Total assembly length was then compared across assemblers and sequencing depths and evaluated against these size estimates to assess concordance with expected genome size. N50 and contig counts were used to assess contiguity, while BUSCO completeness and multi-copy BUSCO percentages were used as proxies for gene recovery and potential duplication artifacts. Base-level accuracy metrics (mismatches and indels per 100 Kbp) were primarily interpreted for the polishing comparison, where differences relative to the unpolished Flye assembly reflected the sequence changes introduced by polishing.

Finally, computational requirements (runtime and maximum RAM usage) for each assembly approach were quantified by accessing SLURM accounting records (command *sacct*). For polishing strategies, runtime was calculated as the sum of the original assembler and the polisher (e.g., Flye = 5min, Racon = 2min, Flye + Racon = 7min), and maximum RAM was taken as the highest value across the runs (e.g., Flye=10GB, Pilon=5GB, Flye + Pilon=10GB).

We summarized read and assembly metrics across fungal genomes using means and their associated 95% confidence intervals, which were used to draw the plots and are reported as mean ± 95% CI.

It is worth noting that two assemblers contributed fewer observations to the computational analyses due to practical execution limits. MaSuRCA failed to complete several assemblies with segmentation faults, an issue that has been reported previously [65], and appears to be dataset- and parameter-dependent. Canu, on the other hand, took over a month to assemble the largest sample at depths higher than 15X, exceeding the available computing infrastructure. Therefore, failed MaSuRCA and incomplete Canu executions were excluded from the analysis.

### Statistical analysis

Statistical analysis was conducted in R v4.4.1 [66] supplemented with tidyverse v2.0 [67] and janitor v2.2.1 [68]. As part of the initial exploratory analysis, we assessed correlations among the response variables using the pairwise Spearman test of the stats::cor() v3.6.2 package on the simulated single-strategy data. Next, assembly performance was analyzed using linear mixed-effects models implemented in lme4 v1.1.37 [69]. Two model classes were fit, distinguished by the depth structure of the assembly strategy. Single-strategy models were used for short-read-only assemblers (SPAdes, ABySS) and long-read-only assemblers (Flye, Hifiasm, Canu, Raven, NextDenovo, Miniasm), with depth modeled as a continuous predictor (*depth_eff*) representing whichever sequencing technology was used. Multi-strategy models were used for hybrid assemblers (SPAdes Hybrid, MaSuRCA) and polishing combinations (E.g., Flye + Polypolish), with separate continuous predictors for long-read depth (*depth_np*) and short-read depth (*depth_il*) and their interaction.

Fixed effects in the single-strategy models included the assembler program, sequencing depth, their interaction, and a genome-characteristic covariate. For simulated data, additional genome features (size, GC content, gene density, and CDS-per-gene ratio) and their interactions with program and depth were included to evaluate how genome architecture modulated assembler performance. For empirical data, no reference genome was available, so the genome-size covariate was computed as the per-isolate mean assembly length across the five Flye assemblies (minimum-overlap settings 1000-3000 bp) at full sequencing depth. This estimate was applied for each isolate as a per-sample constant, and the corresponding full-depth Flye assemblies were excluded from the modeling dataset to avoid using the same observations for both response and covariate estimation. All continuous covariates were centered and scaled prior to fitting, and sample identity was included as a random intercept.

Response variables included contiguity (N50), completeness (BUSCO complete percentage, BUSCO difference from reference), genome recovery (genome fraction recovered, length error percentage), base-level accuracy (mismatches and indels per 100 Kbp), and computational requirements (maximum RAM, runtime). Most responses were transformed prior to modeling to satisfy Gaussian residual assumptions: log10 for responses spanning multiple orders of magnitude (N50, runtime, RAM), logit (with a 0.01 shrinkage offset) for bounded percentage responses near the upper limit (BUSCO complete, genome fraction recovered), and log1p for non-negative rate counts (mismatches and indels per 100 Kbp). Signed differences (BUSCO difference, length error percentage) were modeled on the original scale.

Significance of fixed effects and interactions was assessed using Type III Wald χ² tests with car::Anova v3.1-5 [70] . Post hoc pairwise comparisons among assemblers and sequencing depths were performed with Tukey adjustments using emmeans v1.11.2.8 [71] and multcomp v1.4.28 [72]. Per-program depth slopes and per-program responses to genome covariates were extracted using emtrends. Residual diagnostics were assessed using simulated quantile residuals (n = 1,000) with DHARMa v0.4.7

This allowed us to test whether assembly quality metrics responded differently to increasing sequencing depth across assemblers and whether genome characteristics influenced assembly outcomes. Additionally, we assessed residual diagnostics using simulated quantile residuals (n = 1000) with R package DHARMa v0.4.7 [73], which lead to sensitivity analyses in the single-strategy model by refitting after excluding Miniasm (which produced extreme outlier values for several quality responses), and refitting separately within each strategy (short and long-reads). To enable a direct comparison between the empirical and simulated datasets we predicted values back-transformed to the original response scale. For the simulated dataset, we also derived thresholds from the models to better guide decision process on choosing the depth. Predictions were computed on the transformed response scale, at the mean of all genome-architecture covariates, and at depths 10 to 100x in 5x intervals via lme4::bootMer with 1000 bootstrap replicates to obtain 95% confidence intervals. Thresholds for the decision-guide were defined as the minimum depth at which 90% of the program’s best predicted best value was achieved.

All analyses were run on the Auburn University’s Easley High Performance Computing Cluster using SLURM for job scheduling and resource management. All programs were run in nodes using Intel Xeon Gold 6248R processors with 48 cores and 192 GB of RAM. Jobs were submitted using the *sbatch* command, requesting 10 CPUs and 180 GB of RAM for each assembly. Assemblers were installed using Conda into separate environments, except for ABySS [48] which had its dependencies installed via Conda, and then manually compiled into the environment. Details on the installation and commands used for each program can be found in the GitHub repository.

## Results

### Statistical analyses

Pairwise Spearman correlations among the response variables confirmed that most captured meaningfully distinct components of assembly quality (|ρ| < 0.5, Supplementary Figure S1). The principal exceptions were genome fraction recovered and length error percentage (ρ = 0.86), which both quantify deviation from the reference genome length, and mismatches and indels per 100 Kbp (ρ = 0.79), which both measure base-level errors against the reference.

Initial single-strategy model adequacy assessed using DHARMa simulated residual diagnostics showed dispersion ratios within the expected range (0.95–1.00) across all models, indicating no over- or under-dispersion (Supplementary Table S2). Kolmogorov-Smirnov tests of residual uniformity were statistically significant for all models, as expected given the large per-model sample sizes (n > 5,000), but effect sizes were modest for most responses (KS statistic 0.05-0.18), indicating virtually acceptable fit. BUSCO completeness difference showed the largest deviation (KS = 0.29), due to its strongly skewed distribution in which most assemblies achieved high recovery, even at lower depths.

To confirm that these residual deviations did not affect our conclusions, we refitted the models within each sequencing strategy, where the response distributions were more homogeneous. Similar fixed effects remained significant across all model variants (Supplementary Table S3), and within-strategy models for genome fraction, length error percentage, and mismatches per 100 Kbp showed substantially improved residual uniformity (Supplementary Table S2), confirming that conclusions about assembler-specific performance were robust to residual deviations.

### Simulated data outcomes

The 66 reference fungal genomes we identified via NCBI included a wide array of structural and compositional characteristics (Figure S2). Genome length varied widely (7.4-122.8 Mbp) and was strongly associated with predicted gene content (R^2^ = 0.92, Figure S2A). Although gene content and genome size were strongly associated, this relationship was not completely reflected in the number of coding sequences and introns, as above ∼20 Mbp, the number of predicted CDS and intron features increased more rapidly than the number of genes. This likely reflects the higher compactness of smaller genomes, whereas larger genomes tend to accommodate longer genes, higher intron density, and more complex exon-intron architectures, allowing individual genes to have multiple coding regions [16]. Furthermore, the references ranged from 3 to 23 contigs (average 9.6, Figure S2B), and GC content showed substantial variation among samples, ranging from 28.3% to 62.3% (average 46.7%, Figure S2C).

Across single-technology assemblers, contiguity differed sharply by sequencing technology, with long-read assemblers showing steep N50 gains between 10X and ∼20-30X long-reads depth (LR) before plateauing, whereas short-read assemblers showed minimal response to increasing depth (Figure 1A). For example, Flye increased from 2.72 Mbp (±0.39) at 10X to 3.73 Mbp (±0.77) at 20X and remained stable through 100X. Canu showed an even sharper early rise from 10X to 20X (0.28 Mbp ±0.14 to 3.08 Mbp ±0.65), while SPAdes improved only modestly across depths (0.59 Mbp ±0.12 at 10X to 0.91 Mbp ±0.20 at 100X) and ABySS remained highly fragmented (<0.11 Mbp at all depths). This difference was confirmed by a strong interaction between assembler and sequencing depth (p < 0.001, Figure S3A), consistent with long-read assemblers using read length to establish contiguity rapidly at moderate depth, while short-read assemblers remain limited by repeat-spanning regardless of coverage. From a practical standpoint, long-read contiguity is largely established by 20-40X, and the model-based threshold analysis (Table 1) confirms that Flye and NextDenovo reach 90% of their predicted best N50 by 20X and 10X depth respectively, with further investment offering diminishing returns.

**Figure 1.**
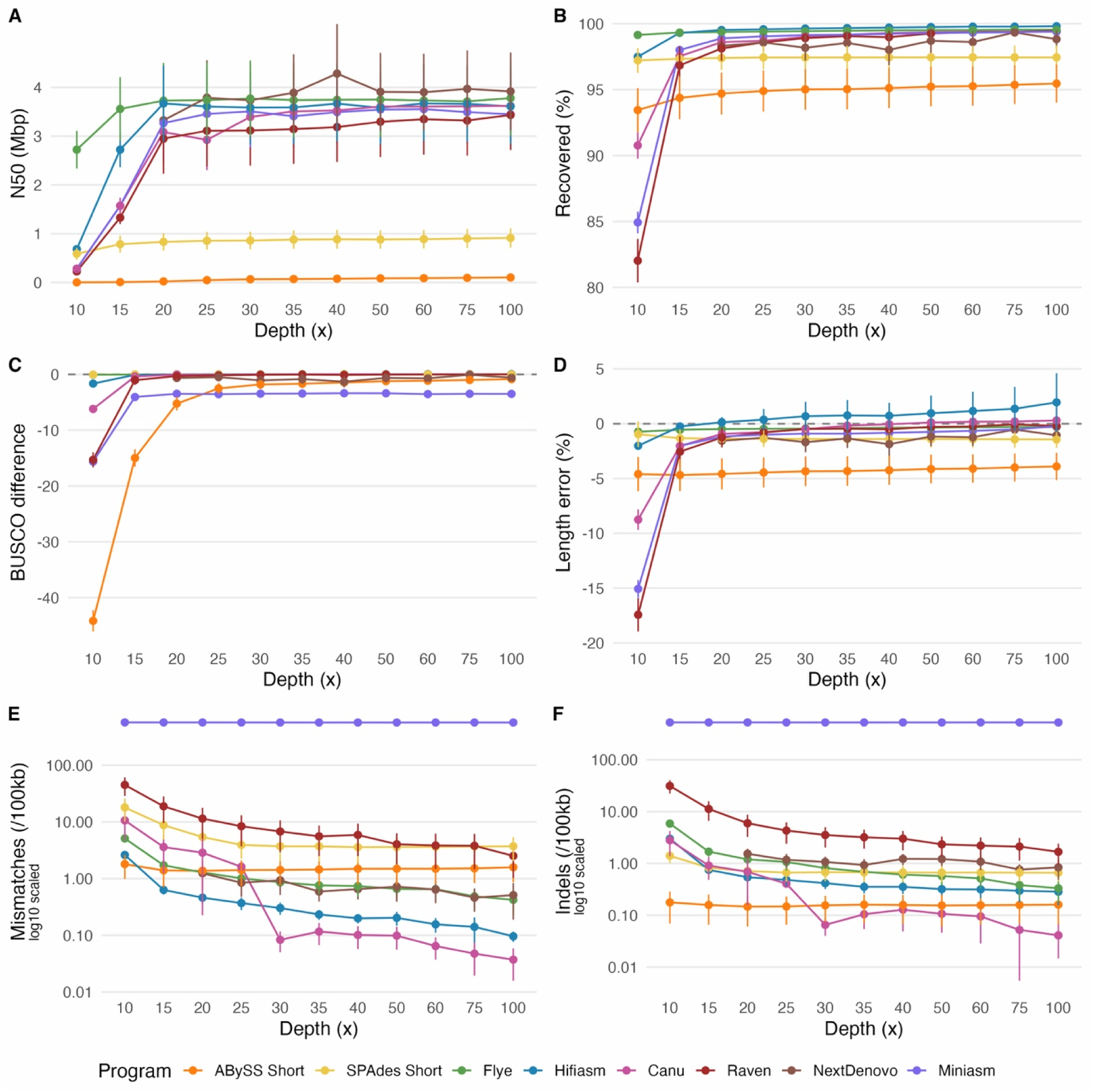
Assembly quality metrics for single assembly strategies. Panels show (A) N50, (B) genome fraction, (C) BUSCO difference, (D) length error, (E) mismatches per 100 Kbp, and (F) indels per 100 Kbp for each assembler as a function of sequencing depth. Error bars represent 95% confidence intervals around the mean. Note the y-axis on panels E & F have been scaled for better visualization. Overall, long-read assemblers showed rapid improvements in contiguity, completeness, and accuracy up to ∼20–40× depth, after which gains plateaued, whereas short-read assemblers exhibited minimal depth-dependent improvement. Alt text (Figure 1): Six-panel figure showing assembly quality metrics for several genome assemblers across increasing sequencing depth. Panels display N50, genome fraction recovered, BUSCO completeness difference from reference, assembly length error, mismatches per 100 Kbp, and indels per 100 Kbp. Long-read assemblers show sharp improvements in contiguity, completeness, and accuracy between low and moderate sequencing depths, while short-read assemblers remain comparatively fragmented with smaller improvements across depths. For most long-read assemblers, metric values stabilize beyond approximately 20-40X coverage.

**Table 1.** Model-based sequencing depth thresholds for each combination of assembler and response variable. Each cell reports the lowest sequencing depth (in X coverage) at which the mixed-effects model first predicts the response to reach within 90% of its best value. Single-strategy programs are reported with its only corresponding value, while multi-strategy programs are reported as “LR | SR” (long-read | short-read depth). Threshold values should be interpreted along with the response values, as low thresholds combined with poor metrics potentially indicate program limitations rather than efficient saturation.

| Metric | N50 | BUSCO completeness | BUSCO difference | Genome recovery | Length error | Mismatches | Indels |
| --- | --- | --- | --- | --- | --- | --- | --- |
| **ABYSS Short** | 100 | 10 | 70 | 10 | 60 | 10 | 10 |
| **Canu** | 95 | 10 | 65 | 10 | 65 | 75 | 80 |
| **Flye** | 20 | 10 | 55 | 10 | 100 | 100 | 95 |
| **Hifiasm** | 85 | 10 | 60 | 10 | 25 | 85 | 100 |
| **Miniasm** | 95 | 10 | 100 | 10 | 70 | 10 | 10 |
| **NextDenovo** | 10 | 10 | 95 | 10 | 95 | 95 | 90 |
| **Raven** | 95 | 10 | 65 | 10 | 65 | 100 | 100 |
| **SPAdes Short** | 65 | 10 | 75 | 10 | 10 | 95 | 85 |
| **ABYSS Hybrid** | 10 \| 100 | 10 \| 10 | 10 \| 70 | 10 \| 10 | 10 \| 65 | 10 \| 10 | 10 \| 10 |
| **Flye + Pilon** | 20 \| 10 | 10 \| 10 | 45 \| 45 | 10 \| 10 | 95 \| 45 | 100 \| 80 | 100 \| 75 |
| **Flye + Polypolish** | 20 \| 10 | 10 \| 10 | 45 \| 80 | 10 \| 10 | 95 \| 30 | 100 \| 10 | 100 \| 10 |
| **Flye + Racon (BWA)** | 20 \| 10 | 10 \| 10 | 55 \| 50 | 10 \| 10 | 95 \| 15 | 100 \| 85 | 100 \| 10 |
| **Flye + Racon (Minimap2)** | 20 \| 10 | 10 \| 10 | 45 \| 80 | 10 \| 10 | 95 \| 10 | 95 \| 100 | 85 \| 95 |
| **MaSuRCA** | 65 \| 100 | 10 \| 10 | 85 \| 15 | 10 \| 10 | 95 \| 10 | 10 \| 100 | 10 \| 100 |
| **SPAdes Hybrid** | 10 \| 75 | 10 \| 10 | 100 \| 95 | 10 \| 10 | 10 \| 10 | 10 \| 95 | 10 \| 85 |

On the other hand, completeness metrics saturated across all assemblers once moderate depth was reached. By 50X, Flye recovered 99.50% (±0.27), Hifiasm 99.75% (±0.17), Raven 99.26% (±0.29), and Canu 99.35% (±0.23) of the reference genome, with differences typically <0.5 percentage points. In contrast, short-read assemblers remained slightly lower even at high depth (SPAdes 97.45% ±0.89; ABySS 95.46% ±1.44 at 100X) (Figure 1B). BUSCO differences relative to the reference similarly stabilized beyond ∼20X for most long-read assemblers (within ±0.1 pp), whereas Miniasm often underperformed (-15.6 pp ±0.93 at 10X and still reduced at high depth) (Figure 1C). Significant “assembler x depth” interaction for BUSCO completeness and genome recovery were found (p < 0.001, Figure S3A), and the threshold analysis confirmed 10X depth to satisfy every assembler tested. These results indicate that sufficient depth is required to approach full completeness, but algorithmic differences become the primary source of variation once moderate coverage is achieved.

Furthermore, base-level accuracy per 100 Kbp (Figure 1E-F) varied widely by the assembler. At 50X, Flye averaged 0.66 mismatches (±0.23) and 0.57 indels (±0.19) per 100 Kbp, whereas Canu achieved even lower values (0.10 ±0.04 mismatches; 0.11 ±0.06 indels) (Figure 1E). Hifiasm similarly showed low mismatch rates (<0.2 per 100 Kbp at ≥40X) but exhibited slight genome length inflation at high coverage (+1.95% ±2.66 at 100X) (Figure 1D). Miniasm stood out by retaining extremely high error rates (>500 mismatches and indels per 100 Kbp at 100X) (Figure 1E-F), explaining its poor BUSCO performance despite moderate contiguity. This stem from the algorithm producing unpolished and uncorrected contig sequences from raw read, in an attempt to reduce computational load [74]. Contrary to expectations, short-read assemblers did not achieve the lowest error rates: at 100X, SPAdes had 3.73 mismatches and 0.66 indels, while ABySS had 1.59 mismatches and 0.16 indels. They also consistently remained under-sized (SPAdes -1.44% ±1.14; ABySS -3.89% ±0.76 at 100X), likely due to collapsing of repeating regions. Assembler choice explained most variation in mismatches and indels, with secondary contributions from depth and their interactions (all p < 0.001, Figure S3A; per-program slopes in Supplementary Figure S4). This suggests that long-read assemblers can achieve comparable or better base accuracy than short-read methods once sufficient depth is reached, while also better preserving the assembly length.

Computational performance varied substantially among assemblers, as maximum RAM and run time grew with depth and reflected differences in the algorithms rather than sequencing technology (Figure 2). For normalized runtime (time per Mbp), the mixed-effects model detected a highly significant “assembler x depth” interaction (p < 0.001), confirming that scaling with depth differed among assemblers (Figure S3A). Additionally, normalizing by genome size drastically reduced the within-assembler variance in runtime, since larger reference genomes incurred proportionally longer runtimes, as confirmed by the significance of the reference size alone in the mixed model for raw time (p < 0.001), reinforcing that genome size is a major driver in resource usage. Normalized memory usage showed strong “assembler x depth” effects (p < 0.001) (Figure 2A, S2A), as at 50X, Flye used 0.32 GB/Mbp (±0.02), Hifiasm 0.66 GB/Mbp (±0.12), Raven 0.35 GB/Mbp (±0.01), and Canu 1.39 GB/Mbp (±0.18). Short-read assemblers were relatively memory-efficient (SPAdes 0.38 GB/Mbp ±0.04 at 100X) but comparatively slow at high coverage (161.2 s/Mbp ±10.8). Canu showed notably higher RAM requirements, and longer runtimes, the latter sufficiently extreme to require separate scaling. At 50X, Flye required 45.1 s/Mbp (±4.06), Hifiasm 30.7 s/Mbp (±8.46), and Raven 13.2 s/Mbp (±0.47), whereas Canu required 1527 s/Mbp (±512) (Figure 2B). Together, these results indicate that while sequencing depth drives improvements in contiguity and completeness up to moderate coverage, computational burden and final assembly quality are largely determined by algorithmic design rather than sequencing technology alone.

**Figure 2.**
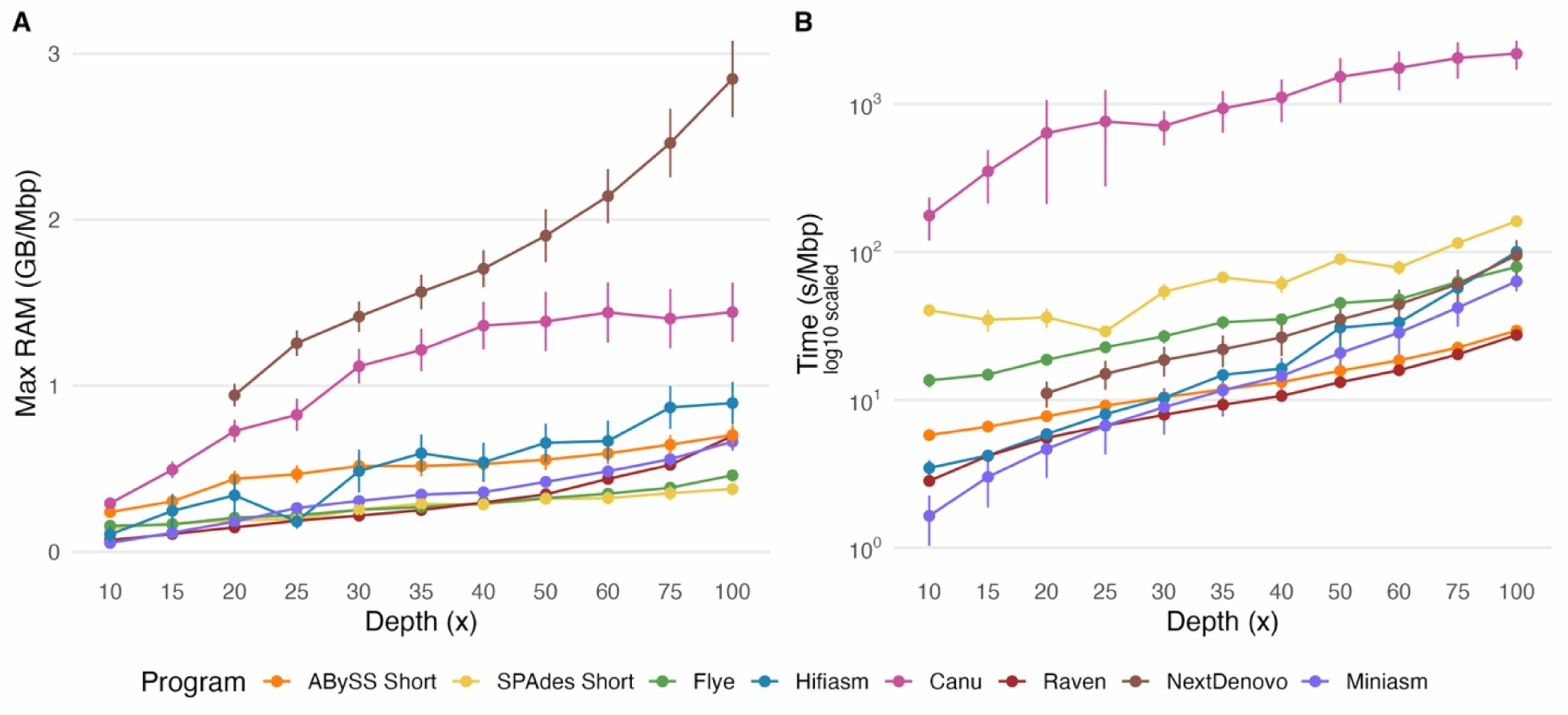
Normalized resource usage per Mbp of assembly for single assembly strategies. Panels show (A) maximum RAM usage per Mbp of assembly and (B) run time per Mbp of assembly for each assembler as a function of sequencing depth. Error bars represent 95% confidence intervals around the mean. Note the y-axis on panel B has been scaled for better visualization. Computational demand increased with sequencing depth but varied strongly by assembler, with Canu and Hifiasm requiring substantially more memory and runtime than other long-read assemblers, while SPAdes remained comparatively slow despite low memory usage. Alt text (Figure 2): Two-panel figure showing normalized computational resource usage per megabase of assembly for different genome assemblers across sequencing depths. Panel A displays maximum RAM usage per mega base pairs, and Panel B shows runtime per mega base pairs. Resource requirements increase with sequencing depth for most assemblers, but the magnitude varies by program, with Canu and Hifiasm showing higher memory and runtime compared to other long-read assemblers, while SPAdes show relatively low memory use but longer runtimes.

Due to Flye’s overall better performance as a stand-alone assembler, we selected it as a representative long-read assembler and added a polishing step with short-reads. This strategy clearly reduced small-variant errors while preserving contiguity (Figure 3). Contiguity (N50) was primarily driven by the long-reads depth (p < 0.001), with no detectable effect of short-reads depth (SR) or polishing program (Figure S5A-B), and genome recovery followed the same pattern. These observations are consistent with the purpose of the polishers to leverage the short-reads to correct at nucleotide-level errors.

**Figure 3.**
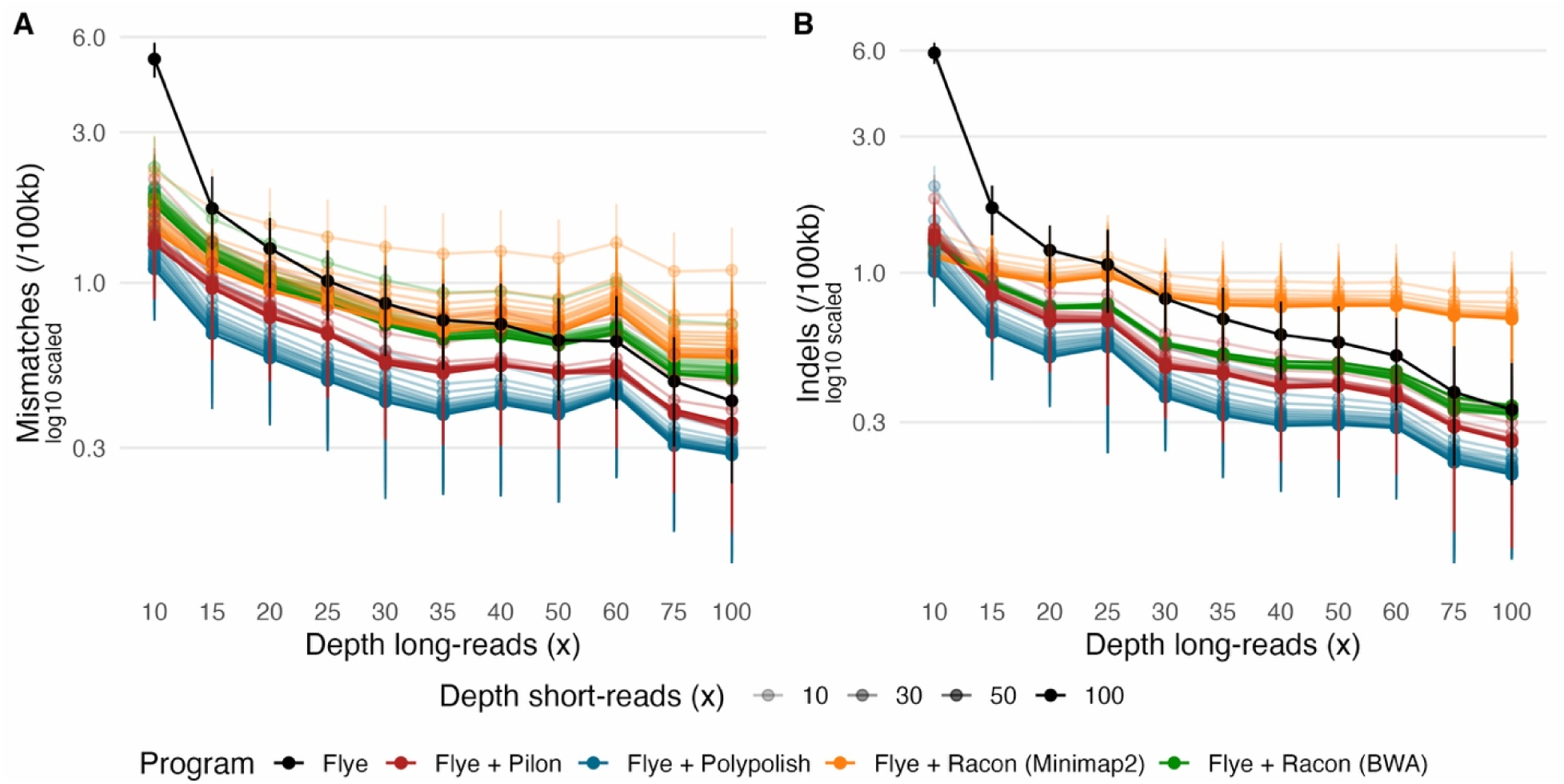
Assembly quality metrics for Flye assemblies before and after short-read polishing. Panels show (A) mismatches per 100Kbp of the original and (B) indels per 100Kbp relative to the original reference genome for the unpolished Flye assembly and Flye assemblies polished with Pilon, Polypolish, or Racon. Racon was evaluated using either Minimap2 or BWA-MEM2 for short-read alignment. Error bars represent 95% confidence intervals around the mean, x-axis indicates long-read depth, and transparency indicates short-read depth. Note the y-axis on all panels have been scaled for better visualization. Polishing noticeably reduced base-level errors, especially at low long-read coverage, with most improvements at moderate short-read depth and lesser gains beyond ∼20-30X. Alt text (Figure 3): Two-panel figure showing base-level error rates in Flye assemblies after polishing. Panel A displays mismatches per 100 kilo base-pairs and Panel B shows indels per 100 kilo base-pairs. The x-axis represents long-read sequencing depth, and point transparency indicates short-read polishing depth. Error rates decrease substantially after polishing, particularly at lower long-read coverage, with improvements stabilizing at moderate short-read depths.

Polishing dramatically reduced base-level errors, particularly when applied to lower-quality long-read assemblies. Short-read depth, long-read depth, and polishing program effects were all highly significant (p < 0.001, Figure S5A-B), while genome architecture covariates had minimal influence on error rates (Figure S5C-F). Using Pilon with 10X SR on the 10X LR unpolished Flye reduced errors to 2.13 mismatches and 1.82 indels per 100 Kbp (±0.53 and ±0.38), whereas Polypolish achieved 1.99 and 2.01 (±0.35 and ±0.28) at the same depths. Increasing short-reads depth to 60X further reduced errors, particularly at low long-reads depth. For example, Polypolish improved from 1.99 mismatches at 10X SR to 1.18 at 60X SR, on the same 10X LR Flye assembly, while at 40X LR + 60X SR reached ∼0.55 mismatches and 0.40 indels per 100 Kbp. However, the magnitude of improvement diminished at higher long-reads depths, indicating that polishing primarily compensates for limited long-read accuracy rather than enhancing already high-quality assemblies. Accordingly, strong base-level accuracy could be achieved at low sequencing cost, provided that Nanopore depth is sufficient to establish contiguity. The model-based threshold analysis (Table 1) also highlighted an eight-fold difference between Polypolish and Pilon in short-read depth requirement in mismatch reduction (10X versus 80X SR, respectively), suggesting that Polypolish is more efficient in extracting the correction from short-reads than Pilon.

Additionally, racon-based polishing showed variable outcomes, as its choice of mapper had a significant impact (Figure 3, Figure S5). Minimap2 outperformed BWA-MEM2 in reducing both mismatches (mean difference = 0.065, p < 0.001) and indels (mean difference = 0.27, p < 0.001). Notably, Racon (Minimap2) was the only combination influenced by genome size (p < 0.05, Figure S5C), evidencing that mapping algorithms differ on how they handle the data. This further demonstrates that each step in the pipeline can significantly influence the final result, even choices that might appear trivial, such as mapper selection.

Since polishing targets corrections at base-level, short-reads impacts on additional quality metrics (N50, length error, genome recovery fraction) were non-significant (Figure S3B & S6), and only marginally altered BUSCO differences . This pattern is consistent with indel correction at lower Nanopore depth improving gene-length integrity and downstream BUSCO alignments.

Polishing added modest computational overhead to the underlying Flye assembly, with both RAM and runtime scaling primarily with short-read depth (Figures 4&5). For Racon-based approaches and Polypolish, memory scaled predictably with short-reads depth from approximately 0.05 GB/Mbp at 10X SR to 0.35-0.42 GB/Mbp at 60X SR (Figure 4A,B,D), reinforcing that read alignment burden increases with short-read depth. Runtime followed similar patterns, with each polisher adding 11-15 s/Mbp at 60X SR. All resource effects were significant for both polishing program and depth (all p < 0.001, Figure S3B, S5A-B) but dominated by the underlying long-read assembler at low short-read coverage and by the polishing step itself when short-read depth exceeded long-read depth (Figures 4&5).

**Figure 4.**
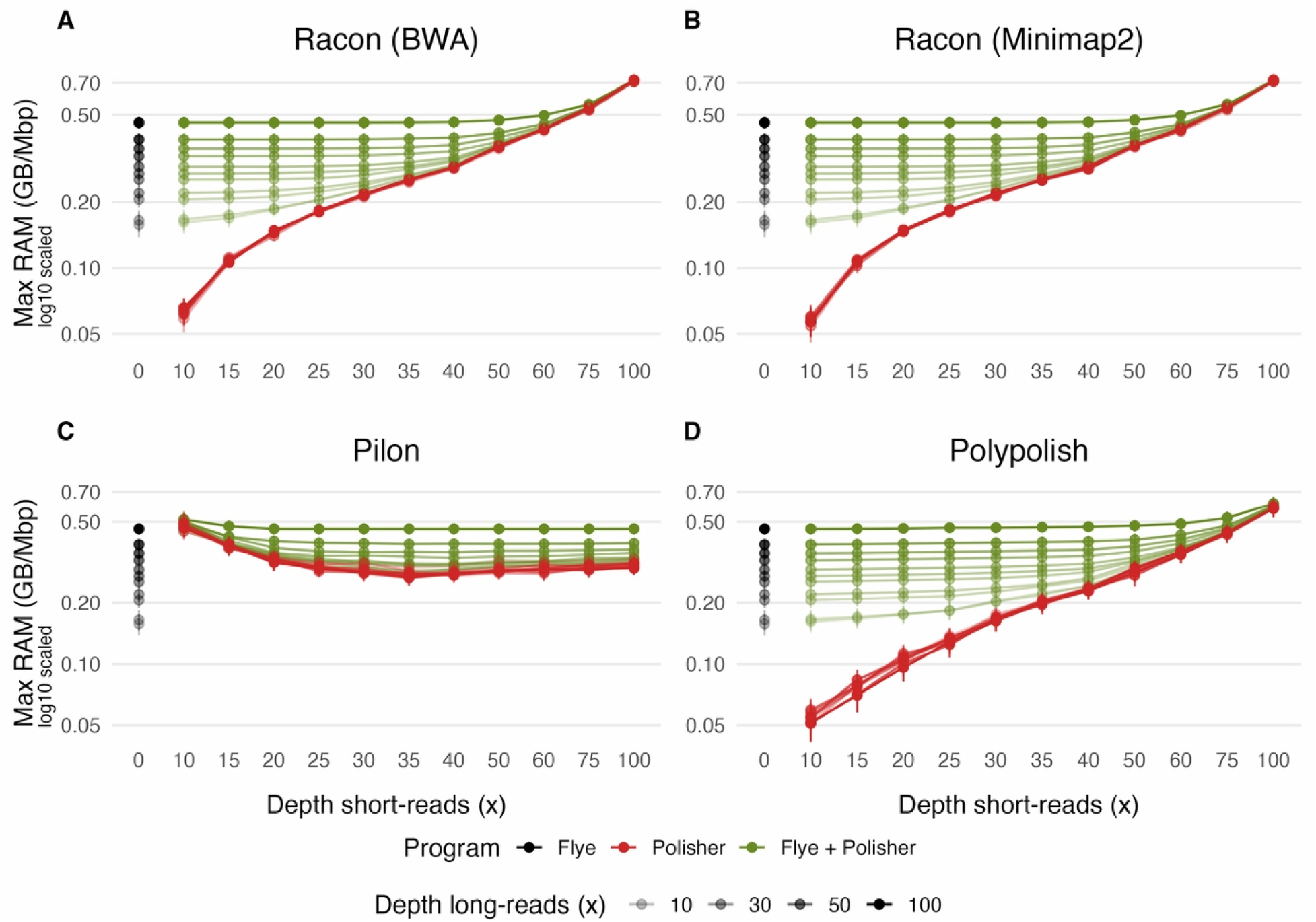
Normalized maximum RAM usage per Mbp for polished assembly strategies. Panels show (A) Racon with BWA-MEM2, (B) Racon with Minimap2, (C) Pilon, and (D) Polypolish. Error bars represent 95% confidence intervals around the mean, x-axis indicates short-read depth, and transparency indicates long-read depth. Note the y-axis on all panels have been scaled for better visualization. RAM usage generally increased with short-read depth for Racon and Polypolish, whereas Pilon showed an inverse pattern with higher memory demand at low coverage, highlighting tool-specific behaviors. Alt text (Figure 4): Four-panel figure showing normalized maximum RAM usage per megabase of assembly for different polishing strategies. Panels correspond to Racon with BWA-MEM2, Racon with Minimap2, Pilon, and Polypolish. The x-axis represents short-read sequencing depth, and point transparency indicates long-read depth. Memory usage generally increases with short-read depth for Racon and Polypolish, while Pilon shows higher memory usage at lower coverage levels.

**Figure 5.**
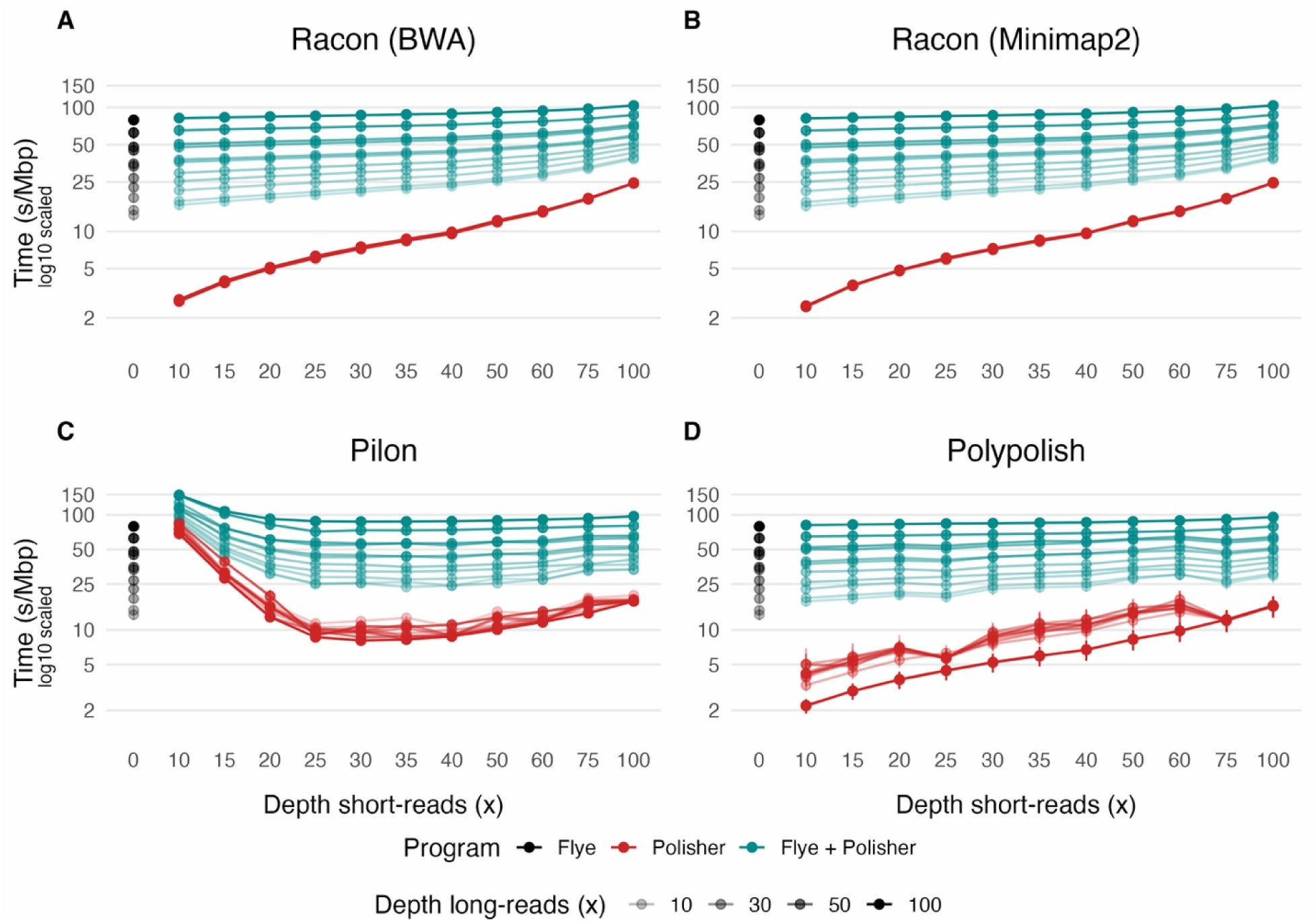
Normalized run time per Mbp for polished assembly strategies. Panels show (A) Racon with BWA-MEM2, (B) Racon with Minimap2, (C) Pilon, and (D) Polypolish. Error bars represent 95% confidence intervals around the mean, x-axis indicates short-read depth, and transparency indicates long-read depth. Note the y-axis on both panels have been scaled for better visualization. Runtime generally increased with short-read depth for Racon and Polypolish, whereas Pilon showed the opposite pattern, with longer runtimes at low coverage that declined and stabilized as coverage increased. Alt text (Figure 5): Four-panel figure showing normalized runtime per megabase of assembly for different polishing strategies. Panels correspond to Racon with BWA-MEM2, Racon with Minimap2, Pilon, and Polypolish. The x-axis represents short-read sequencing depth, and point transparency indicates long-read depth. Runtime generally increases with short-read depth for Racon and Polypolish, while Pilon shows higher runtimes at low coverage that decrease and stabilize as coverage increases.

Unexpectedly, Pilon displayed an opposite pattern (Figure 4C), where RAM usage was higher at low Illumina depth (0.51 [±0.06] GB/Mbp at 10X), then decreased with increasing depth, stabilizing beyond ∼30X (0.33 [±0.03] GB/Mbp). This inverse relationship contrasts with the other polishers and suggests that Pilon’s algorithm may operate less efficiently when coverage is low, possibly due to repeated re-evaluation of ambiguous regions with low support. Consequently, Pilon often required more memory than Flye itself at low short-reads depth. Considering these outcomes, polishing with short-reads reads generally added modest computational overhead to long-read assemblies, with RAM and runtime scaling primarily with short-reads depth, except for Pilon which shows inverse trends likely due to its algorithmic design. Polypolish appeared to offer the best accuracy improvements with moderate resource demands, thus Polypolish-polished Flye assemblies were used as the benchmark long-read baseline for subsequent hybrid comparisons.

Hybrid assembly performance was strongly influenced by algorithm choice. MaSuRCA consistently produced the most contiguous assemblies, reaching 3.53 Mbp (±0.69) Mbp N50 at 30X LR +30X SR, comparable to polished long-read assemblies. SPAdes hybrid achieved intermediate contiguity (∼1.78 Mbp ±0.60), while ABySS Hybrid remained highly fragmented and never exceeded 0.09 Mbp across depths. Increasing short-reads depth beyond ∼30X did not improve contiguity and in MaSuRCA occasionally reduced it slightly, suggesting diminishing returns or conflict between data types at high short-read coverage. These observations were confirmed by the model, which detected significant “assembler x LR” interactions for contiguity (p < 0.001, Figure S3C), indicating that hybrid assemblers differ both in baseline scaffolding capability and in their response to long-read coverage.

Genome recovery (Figure 6B) and BUSCO completeness (Figure 6C) were driven primarily by short-reads depth and assembler choice. MaSuRCA and SPAdes hybrid recovered ≥97.5% of the genome across depths, whereas ABySS Hybrid lagged slightly (95%). BUSCO differences remained small for SPAdes hybrid (∼ -0.2 pp) and MaSuRCA (-0.3 pp), indicating near-complete gene recovery. In contrast, ABySS Hybrid showed severe deficits at low coverage (-44.1 pp ±1.92 at 10X+10X) but improved with short-reads depth, reaching -1.15 pp ±0.54 at 60X SR. These results indicate that short-read support is critical for gene completeness in hybrid assemblies, while long-read depth contributes less once sufficient short-reads coverage is available (p < 0.001 for assembler and “assembler x SR” effects, Figure S3C).

**Figure 6.**
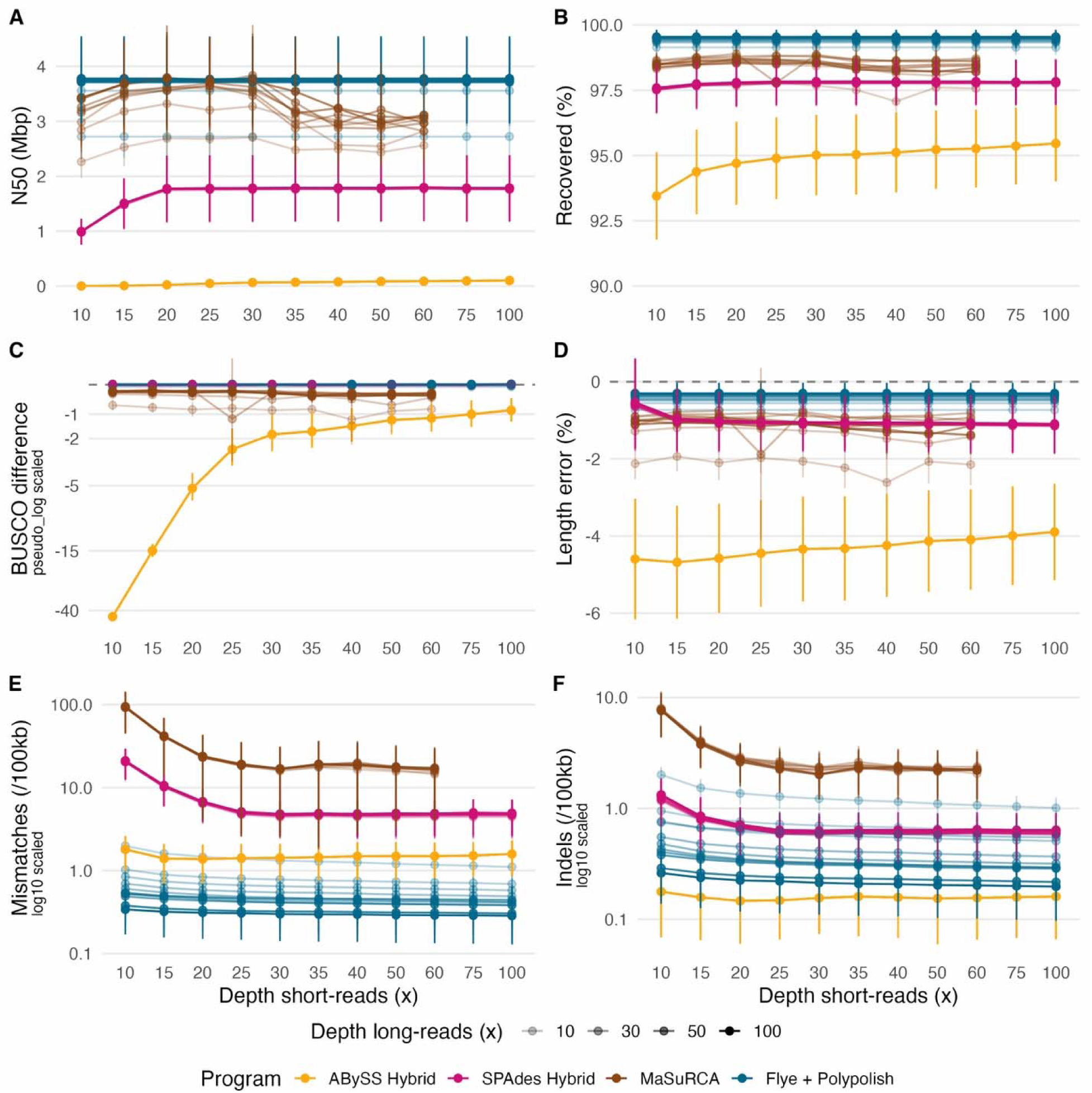
Assembly quality metrics for hybrid assembly strategies. Panels show (A) N50, (B) genome fraction, (C) BUSCO difference, (D) length error, (E) mismatches per 100 Kbp, and (F) indels per 100 Kbp for each assembler as a function of sequencing depth. Error bars represent 95% confidence intervals around the mean, x-axis indicates short-read depth, and transparency indicates long-read depth. Note the y-axis on panels C, E and F have been scaled for better visualization. Hybrid assemblers exhibited clear algorithmic trade-offs: MaSuRCA maximized contiguity, SPAdes hybrid balanced completeness and accuracy, and ABySS hybrid minimized error rates but remained fragmented, while Flye + Polypolish provided a high-quality long-read benchmark across depths. Alt text (Figure 6): Six-panel figure showing assembly quality metrics for hybrid genome assembly strategies across sequencing depths. Panels display N50, genome fraction recovered, BUSCO completeness difference from reference, assembly length error, mismatches per 100 kilo base pairs, and indels per 100 kilo base pairs. The x-axis represents short-read sequencing depth, and point transparency indicates long-read depth. Different assemblers show contrasting performance patterns, with MaSuRCA producing the most contiguous assemblies, SPAdes hybrid showing intermediate contiguity and completeness, and ABySS hybrid showing lower contiguity but reduced base-level error rates; some axes are truncated to improve visualization.

Resource usage reflected these algorithmic differences and scaling behaviors (Figure 7). SPAdes hybrid was the most memory-efficient (∼0.15–0.39 GB/Mbp), while ABySS Hybrid required the most RAM at high depths (up to ∼0.60 GB/Mbp). MaSuRCA memory usage increased primarily with long-reads depth but tended to decrease slightly as short-reads depth increased, suggesting improved read integration efficiency. Runtime scaled steeply with long-reads depth for MaSuRCA and ABySS (e.g., ABySS reaching 179.9 s/Mbp ±192.8 at 60X LR), whereas SPAdes hybrid increased more gradually (∼51 s/Mbp ±4.6 at 10X LR + 60X SR to ∼69 s/Mbp ±9.1 at 60X LR). These patterns indicate that hybrid assembly computational costs are driven largely by long-read data volume and algorithm design rather than short-read depth alone (all p<0.001, Figure S3C, Figure S7).

**Figure 7.**
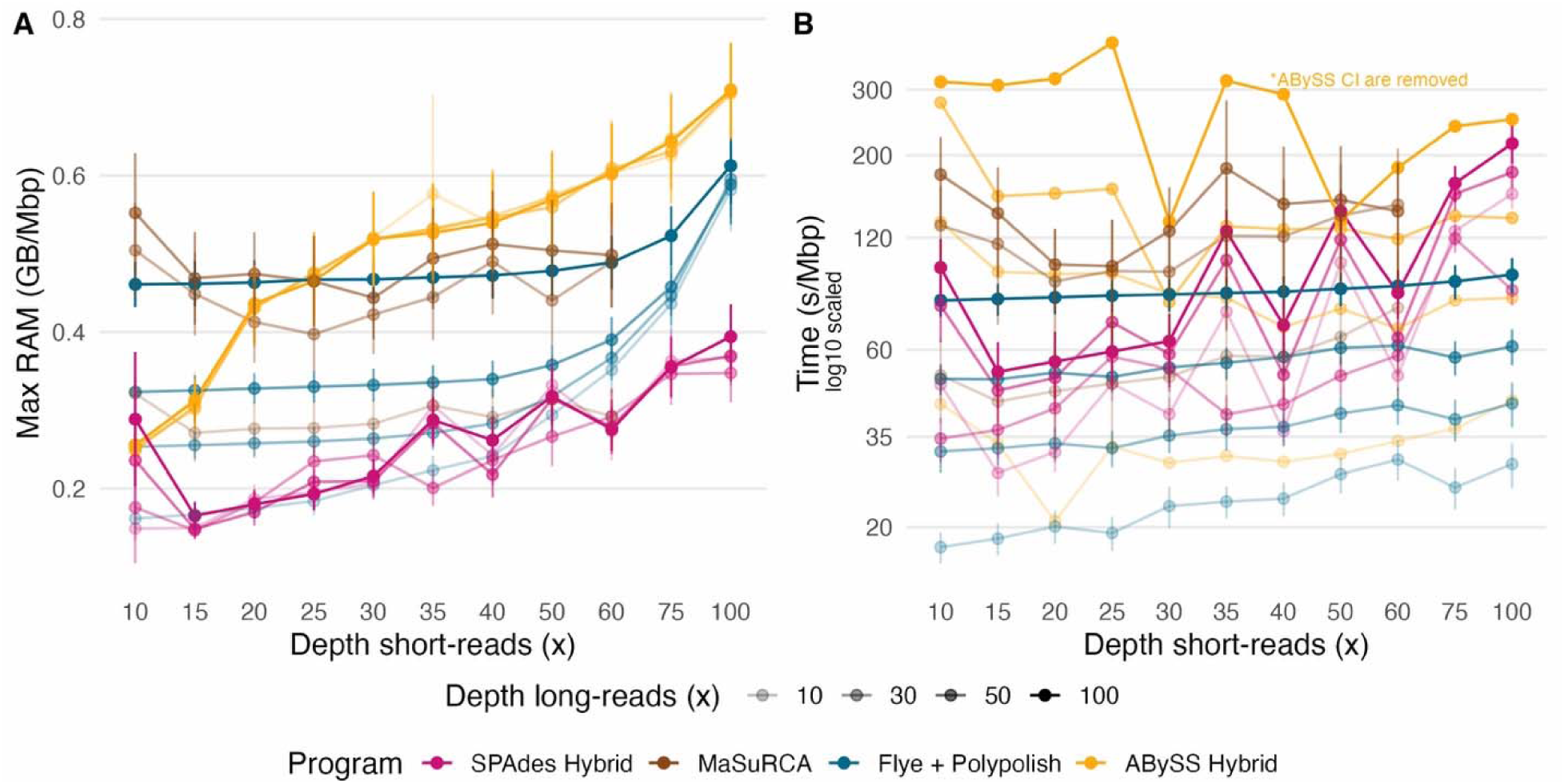
Resource usage per Mbp of assembly for polished and hybrid assembly strategies. Panels show (A) run time per Mbp of reference and (B) maximum RAM usage per Mbp of reference. Error bars represent 95% confidence intervals around the mean, x-axis indicates short-read depth, and transparency indicates long-read depth. Note the y-axis on panel B has been scaled for better visualization, and CI for ABySS Hybrid remove due to high values. Full depth ranges with all CI are provided in Supplementary Figures 8&9 . Computational demand was driven primarily by long-read depth and algorithm design: MaSuRCA and ABySS hybrid showed steep runtime and memory scaling at higher long-read coverage, whereas SPAdes hybrid and Flye + Polypolish remained comparatively resource-efficient and stable across depths. Alt text (Figure 7): Two-panel figure showing normalized computational resource usage per megabase of assembly for polished long-read and hybrid assembly strategies. Panel A displays runtime per mega base pairs, and Panel B shows maximum RAM usage per mega base pairs. Assemblers are compared across sequencing depths, with long-read depth represented in the data points. Resource requirements increase with long-read coverage for some assemblers, particularly MaSuRCA and ABySS hybrid, while SPAdes hybrid and Flye combined with Polypolish show more stable runtime and memory usage across depths.

Overall, hybrid assemblers demonstrate clear algorithmic trade-offs: MaSuRCA maximizes contiguity and completeness, SPAdes hybrid offers balanced performance and efficiency, and ABySS Hybrid prioritizes base accuracy but remains fragmented and underestimates genome size. Despite these strengths, none of the hybrid approaches consistently surpassed polished long-read assemblies, reinforcing that algorithm selection and workflow design remain the primary determinants of assembly quality.

### Empirical data outcomes

Empirical assemblies reproduced the major depth-dependent patterns observed in simulation, although absolute values showed greater variability due to biological heterogeneity and unknown ground truth. K-mer-based genome size estimates closely matched final assembly lengths across assemblers (Figure S10), supporting their use as proxies for genome size. Short-read estimates averaged 36.7 Mb, whereas long-read estimates averaged 22.5 Mb, approximately 38.5% lower. Across assemblies at unsub-sampled depths, final genome lengths typically fell within ∼5-15% of the short reads estimates, indicating good concordance overall. Long-read estimates were typically lower than final assemblies, whereas short-read estimates more closely matched the assemblies. In several isolates, long-read estimates were several megabases smaller than final assemblies, indicating consistent underestimation in those cases. In addition, smaller minimum-overlap parameters in Flye consistently produced longer assemblies, increasing total assembly size by ∼15% on average relative to larger overlap settings, although the change in settings affected the isolates differently, indicating that read-length distribution can also affect final assemblies.

Assembly performance showed significant effects of depth and genome size, as well as “assembler x depth” interactions (all p < 0.001; Figure S11), confirming assembler-specific responses to coverage. Contiguity (N50; Fig. 8A) increased with depth for Flye across all minimum-overlap settings and typically plateaued between 20X and 40X coverage, closely matching the plateau range in the simulated data. Short-read assemblers remained highly fragmented at all depths, reaching similar contiguity only at the unsub-sampled depths (>400X). Assembly size consistently increased with depth (Fig. 8B), with ABySS Short producing the longest assemblies but plateauing at a moderate 25X depth. SPAdes short had the lowest sizes until 35X when it reached similar values to Flye with minimum overlap of 3000 bases. Notably, although SPAdes assembly size was similar to Flye at 35X, they remained highly fragmented, only reaching long-read levels at the ultra-high depths of the unsub-sampled assemblies (>400X).

**Figure 8.**
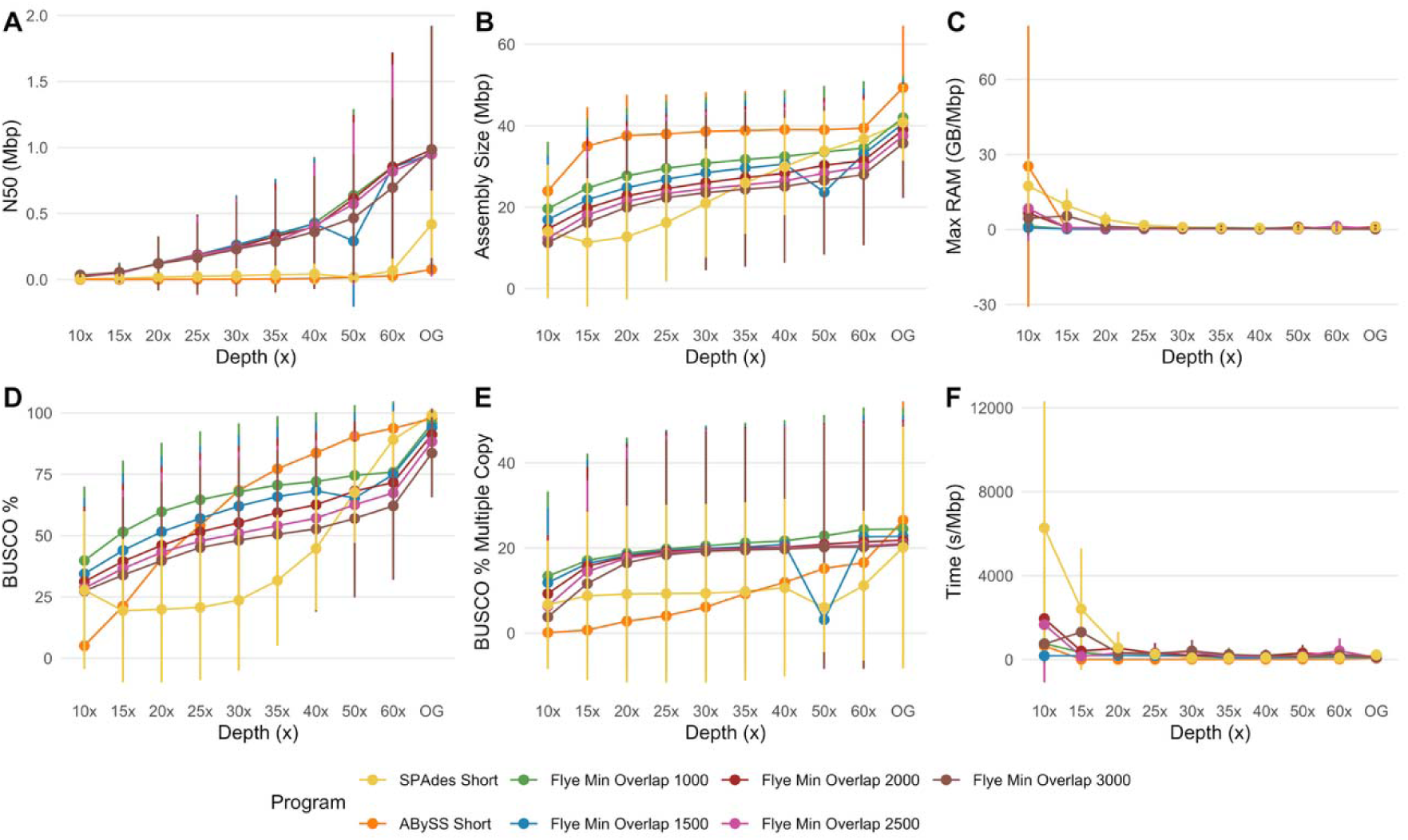
Empirical Data Single Reads Strategy Summary. Panels show (A) N50, (B) Assembly Size, (C) Max RAM per Mbp, (D) BUSCO Complete %, (E) BUSCO Multi Copy %, and (F) Time per Mbp. Error bars represent 95% confidence intervals around the mean. Across empirical isolates, increasing long-read depth improved contiguity and completeness while reducing computational cost, with Flye outperforming short-read assemblers and minimum-overlap settings influencing assembly size and recovery. Alt text (Figure 8): Six-panel figure summarizing assembly metrics for empirical fungal genome assemblies generated using single-read strategies. Panels display N50, total assembly size, maximum RAM usage per megabase, percentage of complete BUSCO genes, percentage of multi-copy BUSCO genes, and runtime per megabase. Error bars indicate 95% confidence intervals around the mean values across isolates. Increasing long-read sequencing depth generally corresponds with higher assembly contiguity and completeness, while short-read assemblers show lower contiguity and similar or greater computational demands.

Completeness improved noticeably with depth (Fig. 8D), as depth and genome size significantly influenced BUSCO completeness (p < 0.001; Figure S11 and S12). Flye assemblies exceeded ∼70% BUSCO completeness at moderate coverage (≥25X) under flexible overlap settings, while short-read assemblies diverged from one another. ABySS recovered more BUSCO genes than SPAdes and at comparable levels to Flye, while SPAdes required at least 50X to achieve similar results. Furthermore, multi-copy BUSCO percentages remained low overall (Fig. 8E), though variance increased with depth for the long-reads, while SPAdes and ABySS remained unchanged until the unsubsampled depths, suggesting a stronger but limited duplication artifacts with long-reads, but strong isolate-specific effects.

Assembly size accuracy and computational demand also stabilized with increasing depth. Unlike the simulated datasets, computational requirements were highest at low coverage and declined rapidly as depth increased, particularly for SPAdes, with both memory (Fig. 8C) and runtime (Fig. 8F) stabilizing beyond ∼25-30X. Mixed-effects models confirmed significant assembler, depth and genome size effects on runtime and memory use (p < 0.05; Figure S11 and S12), indicating that incomplete assemblies at low depth impose greater computational overhead, possibly due to fragmented graph structures and ambiguous paths. Finally, comparison of polished versus unpolished assemblies (Figs. S12C-E & S13) showed that polishing primarily improved low-depth assemblies. Base-level corrections were greatest at low long-read coverage and increased sharply with initial short-read depth before plateauing near ∼20-25X, consistent with simulation results.

Across all measured outcomes, empirical assemblies largely followed the qualitative patterns observed in simulation (Fig. 9), confirming that the inferences derived from simulated data translate to field-collected fungal isolates. Most discrepancies were quantitative rather than qualitative: empirical Flye N50 continued to gain through 60X long-read coverage where simulated Flye N50 plateaued by 20-30X, and computational requirements at low depth were probaby inflated by fragmented graph structures in empirical but not in simulated assemblies. These differences suggest that simulation-derived depth recommendations should be taken as lower bounds for real-world sequencing planning, with empirical fungal assembly likely requiring approximately 1.5-2X the simulated depth to achieve equivalent contiguity.

**Figure 9.**
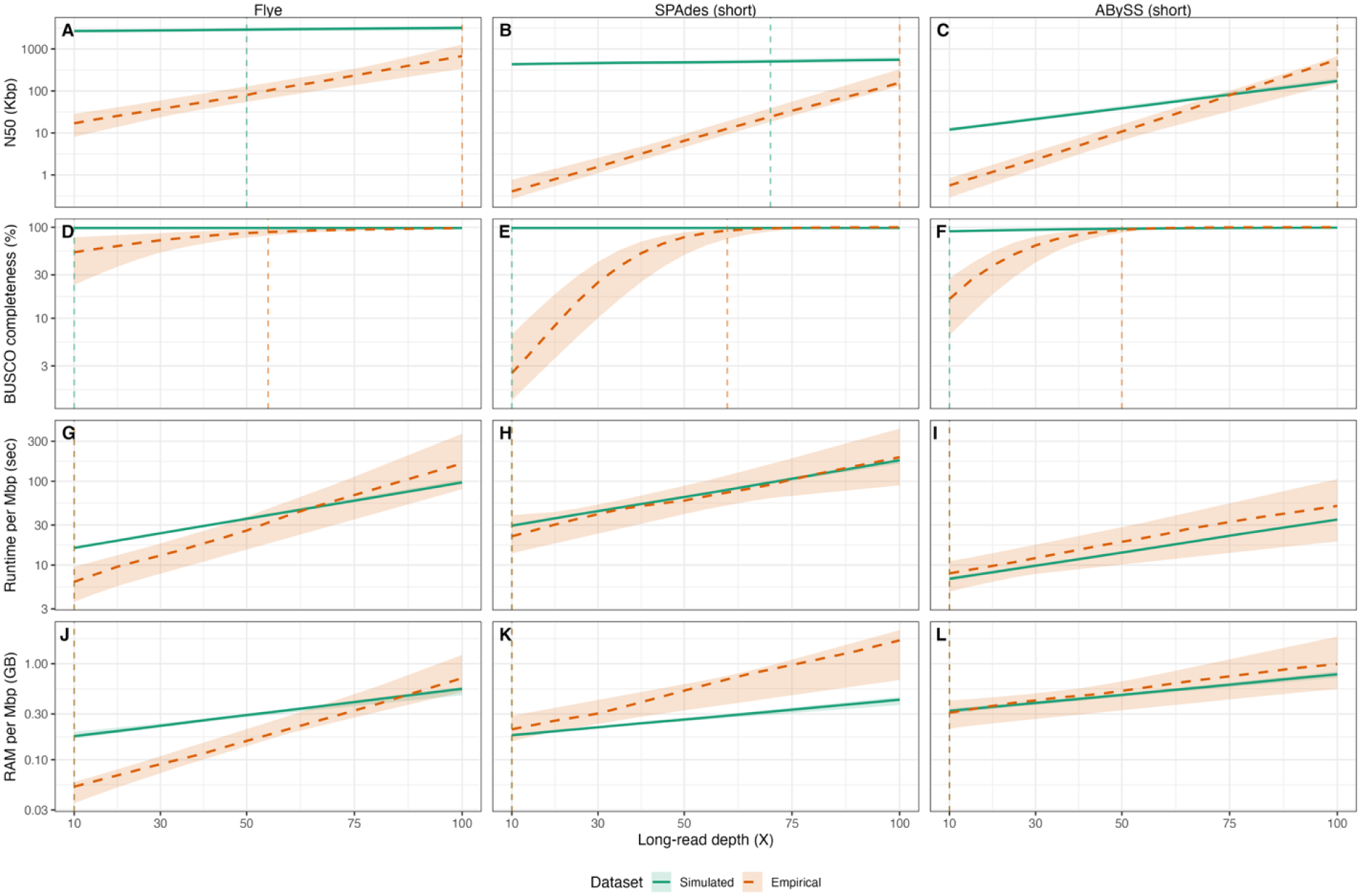
Model-predicted depth-response curves for shared assemblers and metrics across simulated and empirical datasets. Rows represent the assembly metrics: (A, B, C) N50; (D, E, F) BUSCO Completeness; (G, H, I) run time; (J, K, L) max RAM. Columns represent different programs: (A, D, G, J) Flye; (B, E, H, K) SPAdes Short; (C, F, I, L) ABySS Short. Orange-dashed lines represent the empirical dataset and green-continuous lines represent the simulated dataset. Shaded ribbons represent 95% confidence intervals. Vertical line of each color represents the depth threshold at which 90% of its best value was achieved. All y-axes use log10 scaling. Simulated predictions for runtime and RAM use reference-normalized metrics, while empirical predictions use proxy-normalized. Predictions were truncated at biological lower bounds where applicable (e.g., non-negative for runtime and RAM). Alt text (Figure 9): Twelve line-plots in four rows (N50, BUSCO completeness, runtime per megabase, RAM per megabase) by three columns (Flye, SPAdes short-read, ABySS short-read), plotting model predictions against long-read depth (10 to 100X) on log10 y-axes. Green solid lines show simulated data, orange dashed lines show empirical data, with 95% confidence ribbons and vertical depth-threshold markers. Flye achieves the highest N50, with simulated curves plateauing earlier and higher than empirical. BUSCO completeness displays a sharp curve on the short-reads assemblers, whereas Flye manages much higher completeness at lower depths. Runtime and RAM requirements trends upwards with increased depth at a relatively constant rate and does not appear to saturate.

## Discussion

This study addresses a gap in genome-assembly by focusing on fungi whose genome architectures (broad size range, variable gene structure, and frequent repeats) pose different challenges from the bacterial benchmarks that dominate much of the assembly literature [36–38,42]. Using a combined framework of simulated and empirical datasets spanning diverse genome architectures, our results showed that assembly outcomes reflect a three-way interplay between sequencing depth, algorithmic design, and intrinsic genome properties. In particular, depth effects were strongly assembler-dependent, with long-read assemblers showing larger contiguity gains at low-to-moderate long-reads coverage followed by plateauing, while short-reads-only assemblers showed comparatively weak responses to increased depth.

These contrasting responses reflect fundamental differences in assembly paradigms: long-read assemblers leverage read length to span repeats and resolve structural complexity once sufficient overlap coverage is achieved, while short-read assemblers rely on de Bruijn graph connectivity, which remains constrained by repeat length and sequencing errors even at very high coverage. This pattern supports a practical interpretation of fungal assembly performance: additional data does not improve all approaches equally, and sequencing depth must be evaluated alongside assembler error tolerance, repeat-resolution capacity, and graph construction strategy.

Across approaches, our results point to diminishing returns in long-read depth once moderate coverage is achieved. In both simulated and empirical datasets, most improvements in contiguity and completeness occurred by ∼20-40X long-read depth (Figure 9), after which additional data produced only marginal improvements despite increased computational cost. Similar saturation patterns have been reported in long-read benchmarking studies, where assembly contiguity and completeness improve rapidly at low-to-moderate coverage but plateau once repeat-spanning coverage and graph connectivity are achieved [37,38,75,76]. Importantly, this “coverage-to-benefit” curve differed among assemblers, reinforcing that “optimal depth” is not a single number but depends on the algorithm’s sensitivity to read errors, repeat structure, and coverage heterogeneity.

Furthermore, short-read polishing presents an efficient alternative to pushing long-read depth beyond this moderate range. In the simulated polishing analyses (Flye baseline), short-read depth and polisher choice strongly shaped mismatch/indel outcomes, while contiguity and genome recovery were largely inherited from the underlying long-read assembly(Figs. 3, 4, 5, S3B, S5). Similarly, in the empirical comparisons, polishing base corrections were higher in lower-depth long-read assemblies, with sharp increased gains at lower short-read depths before plateauing (Fig. S10). These results reinforce prior reports that polishing can deliver large base-level accuracy improvements at relatively modest additional sequencing cost, often approaching maximal benefit at moderate short-read depths [32,77]. Among polishers, Polypolish consistently provided the strongest improvements, while Racon outcomes strongly depended on mapper choice, emphasizing that seemingly “upstream” decisions can propagate to measurable differences in final accuracy, a point often emphasized in workflow comparisons in genomics [78–80].

Hybrid assembly results further illustrate how algorithm choice shapes outcomes even when the data types are fixed. Hybrid assemblers showed greater variability than other assembler types, reflecting differences in how each assembler balances short- and long-read information, and resulting in clear trade-offs in contiguity, accuracy, and computational cost (Figs. 6&7). MaSuRCA, tended to maximize contiguity, consistent with its strategy of constructing super-reads and then leveraging long reads for scaffolding/bridging [53], but this came with higher error rates and heavier computational demands in our benchmarks. In contrast, hybridSPAdes generally produced less contiguous but cleaner assemblies, consistent with a short-reads-first-assembly backbone with cautious long-read integration to fill gaps and resolve repeats [33], thereby trading contiguity for improved base-level accuracy. By comparison, ABySS hybrid attempts to rescaffold short-reads draft contigs using long-reads data [48], resulting in high base-level accuracy but highly fragmented assemblies, suggesting that its strategy may under-utilize long-read information relative to other hybrid approaches. Overall, our data suggest that algorithm selection can meaningfully shape assembly structure and error profiles even when sequencing depth and data composition are held constant, reinforcing the importance of evaluating multiple workflows rather than relying on a single default pipeline.

Genome-to-genome variation further indicates that assembly success is not uniform across fungal taxa, and that genome characteristics can modulate both assembly outcomes and computational requirements. In the simulated data, genome size and gene density were consistently associated with greater variability in contiguity, completeness, and computational demand (Figs. 4C-D, 5D-E, 7D-E) indicating that larger and more structurally complex genomes require proportionally greater sequencing and computational resources. These relationships help explain part of the dispersion observed in the empirical datasets and further suggests that there is no single best solution, and that recommendations should be scaled with genome complexity. Although our models do not include a dedicated repeat-content term, repeat and transposable-element content is a known driver of assembly-graph ambiguity, particularly for short-read approaches [81,82]. In fungal genomes, these are known to be drivers of genome size and gene density [25,83,84], both of which are explicit covariates in our analysis. As such, the genome-architecture effects we show, including the higher computational demand and greater depth requirements for larger genomes, already encompass much of the influence that could be attributed to repeat content. Structurally compact genomes achieved near-maximal performance at lower coverage while larger, intron-rich, or repeat-dense genomes benefited more from long-read depth and polishing. Despite base composition varying widely across the sampled fungi, its influence on final assembly quality was comparatively limited relative to structural features, suggesting that repeat organization and gene architecture, rather than GC content alone, are the primary determinants of assembly difficulty.

Patterns observed under controlled simulation were quantitatively recapitulated in the empirical datasets (Fig. 9), indicating that simulation-based benchmarking can meaningfully anticipate real-world assembly behavior despite increased biological variability and lack of a known ground truth. Depth thresholds for contiguity and completeness stabilization differed by approximately 10-20% between simulated and empirical assemblies, and the relative performance ranking of major long-read assemblers remained consistent across both datasets. Polishing gains likewise plateaued within comparable short-read coverage windows (∼20–25X in simulation vs. ∼20X in the empirical data), demonstrating that depth-dependent trade-offs identified in simulation translated closely to practical sequencing conditions. Although absolute metric values varied among isolates due to genome-specific features and laboratory variability, the agreement in performance trends indicates that controlled simulation can serve as a practical planning heuristic for fungal genome assembly. This concordance aligns with broader benchmarking efforts in genomics that emphasize the utility of simulated datasets for evaluating algorithmic behavior [39,42]. Our results extend this conclusion to eukaryotic microbial genomes where structural complexity and repeat content introduce additional assembly challenges.

Overall, our data suggest that laboratories can optimize their sequencing strategies by targeting moderate depths (e.g., 30-40X long-read + 10-20X short-read) to achieve high-quality assemblies with manageable costs and computational burdens. Platform selection, however, is often shaped by local infrastructure, budget, and experimental scale. Short-read technologies remain highly standardized with lower per-base costs and high base-level accuracy, but often depend on centralized instrumentation or external core facilities to realize their throughput advantages [85,86]. In contrast, long-read platforms such as the MinION often require lower initial capital investment and offer greater portability and flexibility for in-house or field-based sequencing, making them attractive options (Table 2). Currently, the newest Oxford Nanopore Technologies MK1D platform is priced around $5,000 (reagents and flow cells included), while short-reads’ entry-level iSeq 100 ($20,000) has been phased-out, making the MiSeq i100 ($50,000) the cheapest option. Furthermore, this Nanopore option is much more portable, allowing for flexible deployment in both laboratory and field settings [87]. For fungal genomes, which are typically smaller than mammalian or plant genomes, the cost difference becomes less pronounced, although laboratory effort can also differ. Short-reads tolerates fragmented DNA, simplifying extraction protocols, whereas long-reads sequencing typically requires high-molecular-weight DNA with higher quality to maximize read length. This makes a significant difference for fungi, whose tough cell walls and presence of secondary metabolites can complicate DNA recovery [88–90]. Although combining short- and long-read data consistently produced the most reliable assemblies in this study, dual-platform workflows increase library preparation effort and overall cost and may not be necessary for all projects. Instead, our results indicate that carefully balancing sequencing depth, platform choice, and algorithm selection can often achieve comparable assembly quality without excessive investment in raw data volume alone.

**Table 2.**
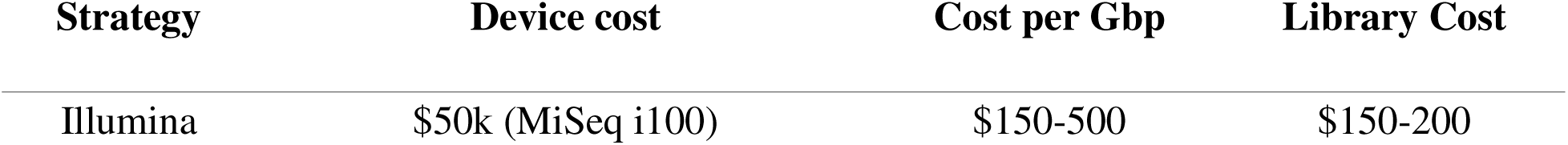
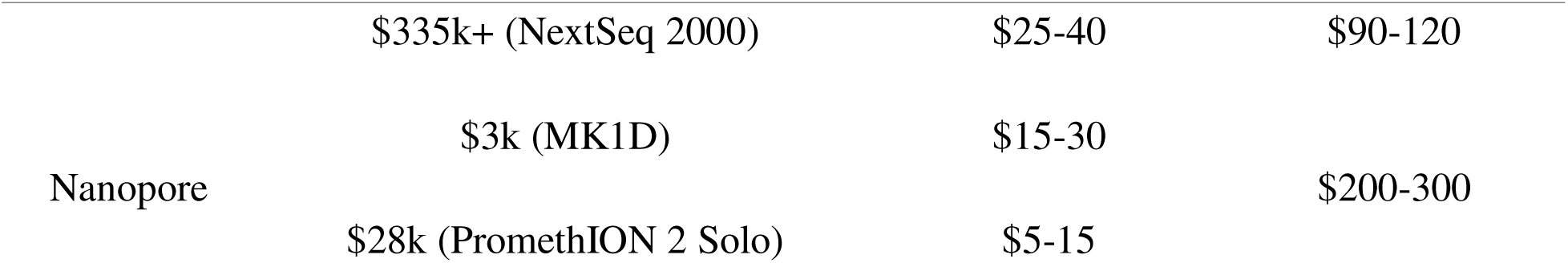
Comparison of practical considerations across sequencing and assembly strategies. Costs are approximate 2025 market estimates based on publicly available provider pricing and typical commercial sequencing service rates, rather than instrument purchase costs alone. Depending on the provider, these costs may include consumables, library preparation, sequencing reagents, instrument operation, and service overhead.

## Conclusions

This study demonstrates that, for fungal genomes, sequencing depth alone does not determine assembly quality; algorithm choice and genome architecture exert an equal or greater influence on assembly quality. Long-reads sequencing delivered the biggest gains in contiguity at moderate depths (∼20-30X), and adding modest short-read-based polishing (∼10-20X) was usually sufficient to achieve high base accuracy without substantial increases in cost or computational demand. Assembly performance varied with genome size and gene density, which were associated with differences in both contiguity and computational demands, although GC content played a comparatively minor role relative to structural features. Together, these results indicate that effective assembly design should balance software selection with the biological complexity of the target genome rather than relying on sequencing depth alone. For most fungal genomes, ∼30-40X long-read depth combined with ∼10-20X short-read polishing provides an efficient compromise between quality, cost, and computational burden. Within this framework, the Flye + Polypolish workflow showed the most consistent performance across simulated and empirical datasets, outperforming alternative polishers and many hybrid approaches overall. Nevertheless, workflow selection can be tuned to match available computational resources, genome complexity, and project priorities, and recommendations should remain adaptable as new assemblers and polishing strategies emerge.

## Acknowledgements

We thank members of the Willoughby Lab for constructive comments on earlier versions of this work. In addition, we are grateful to the many field assistants that worked collecting the original fungal samples used in our sequencing efforts. We also thank Mick Persyn, Samantha Huey and Tess Lindow who assisted with fungal culturing and DNA extractions. This work was completed in part with resources provided by the Auburn University Easley Cluster.

## Author Contributions

G.A.A.S. and J.R.W. conceptualized, planned the study design and developed the benchmarking framework. Empirical data, including sample collection, culturing, and DNA extractions and sequencing were done by G.A.A.S., T.R.F., and L.G.E. Supervision and resource acquisition were done by J.R.W. and L.G.E. The original draft was written by G.A.A.S., and subsequent revisions and editing were done by all authors. All authors read and approved the final manuscript.

## Supplementary Files

Supplementary table 01:

File name: Silva_benchmarking_supp-tables.xlsx

File format: xlsx

Title: Supplementary Table 01

Description: Record of assemblies downloaded from NCBI RefSeq and used in the simulated part of the study

Supplementary table 02:

File name: Silva_benchmarking_supp-tables.xlsx

File format: xlsx

Title: Supplementary Table 02

Description: DHARMa residual diagnostics for all simulated mixed-effects models, reporting the Kolmogorov-Smirnov statistic and p-value, dispersion ratio and p-value, and outlier-test p-value for each response variable and model variant, along with the response transform and observation count.

Supplementary table 03:

File name: Silva_benchmarking_supp-tables.xlsx

File format: xlsx

Title: Supplementary Table 03

Description: Fixed-effect coefficient estimates for all simulated mixed-effects models, reporting each coefficient with its standard error, 95% confidence interval, test statistic, and p-value, on the transformed response scale, for the single-strategy and multi-strategy frameworks.

Supplementary table 04:

File name: Silva_benchmarking_supp-tables.xlsx

File format: xlsx

Title: Supplementary Table 04

Description: Type III Wald chi-square omnibus tests for each fixed-effect term in the simulated mixed-effects models, reporting the chi-square statistic, degrees of freedom, and p-value for every term across all response variables and model variants.

Supplementary table 05:

File name: Silva_benchmarking_supp-tables.xlsx

File format: xlsx

Title: Supplementary Table 05

Description: Per-program depth-response and genome-covariate slopes for the simulated mixed-effects models, estimated via emmeans::emtrends, reporting each slope with its 95% confidence interval and p-value on the transformed response scale.

Supplementary table 06:

File name: Silva_benchmarking_supp-tables.xlsx

File format: xlsx

Title: Supplementary Table 06

Description: Fixed-effect coefficient estimates for all empirical mixed-effects models, reporting each coefficient with its standard error, 95% confidence interval, test statistic, and p-value, on the transformed response scale, for the single-strategy and multi-strategy frameworks.

Supplementary table 07:

File name: Silva_benchmarking_supp-tables.xlsx

File format: xlsx

Title: Supplementary Table 07

Description: Type III Wald chi-square omnibus tests for each fixed-effect term in the empirical mixed-effects models, reporting the chi-square statistic, degrees of freedom, and p-value for every term across all response variables and model variants.

Supplementary table 08:

File name: Silva_benchmarking_supp-tables.xlsx

File format: xlsx

Title: Supplementary Table 08

Description: Per-program depth-response and genome-size slopes for the empirical mixed-effects models, estimated via emmeans::emtrends, reporting each slope with its 95% confidence interval and p-value on the transformed response scale.

Supplementary figures:

File name: Silva_fungi_assembly_benchmarking_supplementary-figures.docx

File format: docx

Title: Supplementary Figures

Description: Compilation of all supplementary figures mentioned in this manuscript.

