## Supplemental Figures for "Quantitative comparison of fungal genome assembly strategies using short and long-reads from simulated and empirical sequencing data"

**Supplementary Figures**

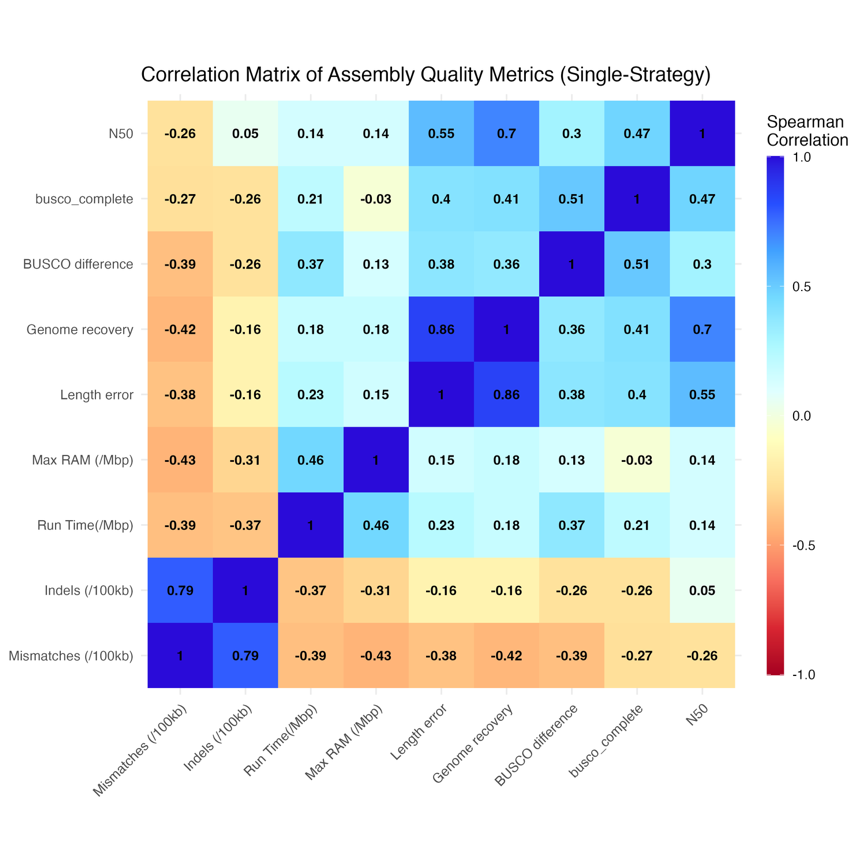

Supplemental Figure 1. Pairwise Spearman correlations between response variables for single-strategy simulated assemblies. Cell color indicates correlation strength and direction (blue, positive; white, zero; red, negative), with ρ displayed in each cell. Note that negative correlations between "higher-is-better" and "lower-is-better" metrics reflect their inverse correlation rather than independence. Most responses were weakly correlated (|ρ| < 0.5), indicating that they capture distinct components of assembly quality, and the strongest were between genome fraction recovered and length error percentage (ρ = 0.86) and between mismatches and indels per 100 Kbp (ρ = 0.79).

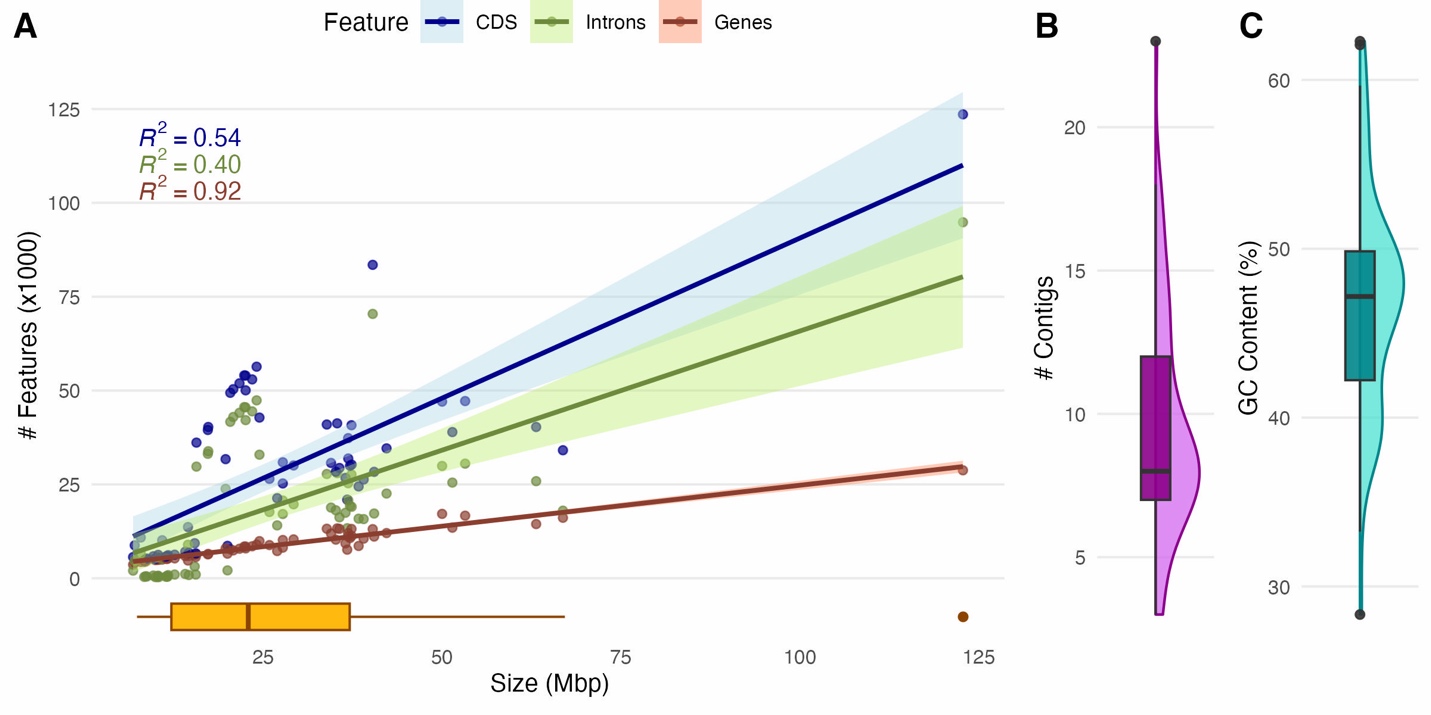

Supplemental Figure 2. Characteristics of the reference genomes used in the assembly benchmark. (A) Number of genes, coding sequences (CDS), and introns versus genome size (in box-plot). (B) Distribution of contig counts. (C) Distribution of GC content. These genomes cover broad diversity, with genome size strongly associated with gene content and intron complexity, providing a representative range of fungal genomic architectures for benchmarking.

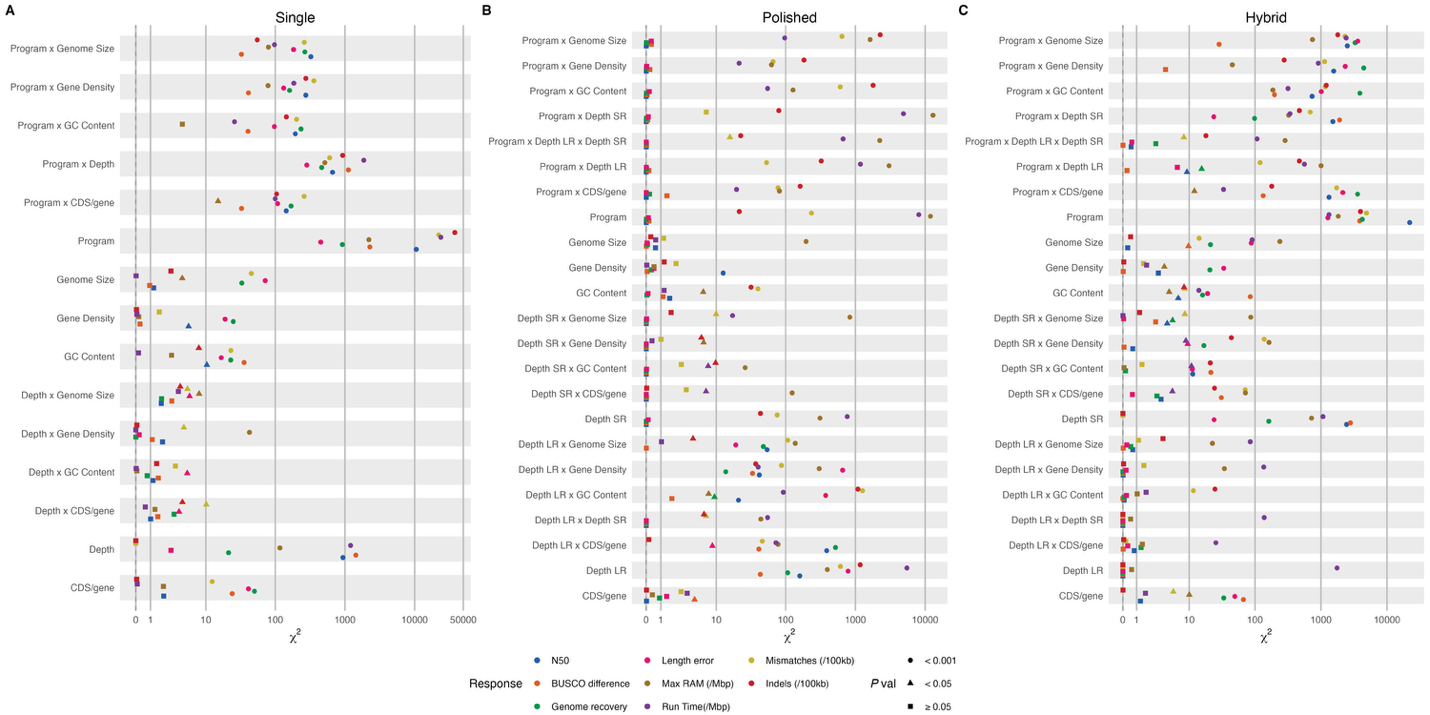

Supplemental Figure 3. Forest plot of Type III Wald χ² statistics for all fixed-effect terms across simulated mixed-effects model. Panels indicate sequence strategy: (A) single sequencing, (B) polishing, (C) hybrid sequencing. Color indicates the response: N50, BUSCO completeness, BUSCO completeness difference from reference, genome fraction recovered, length error percentage, mismatches and indels per 100 Kbp, RAM per Mbp of reference genome, and runtime per Mbp of reference genome. Shape indicates the significance level: square p ≥ 0.05, triangle p < 0.05, circle p < 0.001. Points represent the χ² statistic for one fixed-effect term in one response model, plotted on the pseudo-log scaled x-axis.

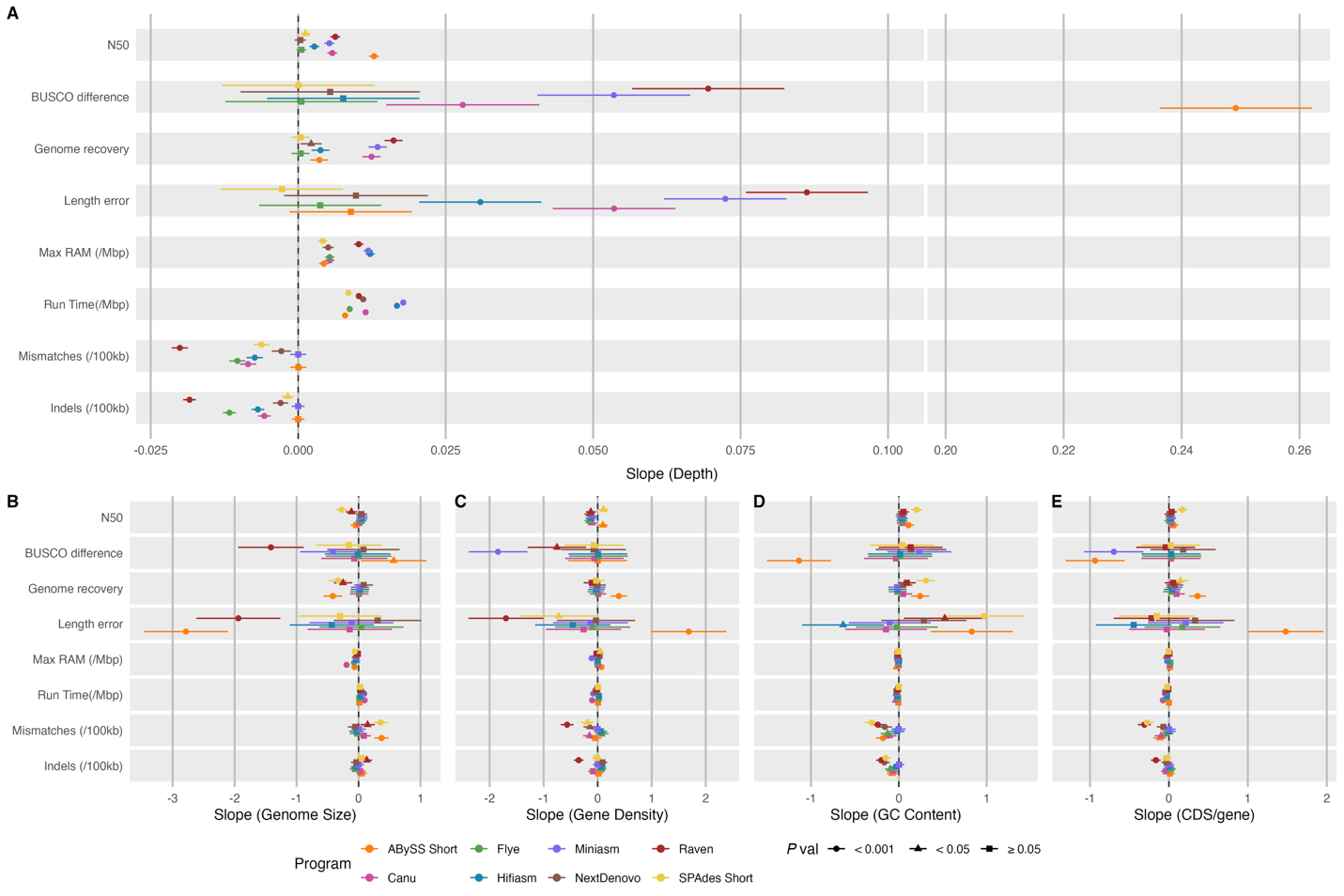

Supplemental Figure 4. Forest plot of per-program slopes for continuous predictors in the simulated single-strategy model. Slopes were estimated via emmeans::emtrends, with points representing the per-program slope and horizontal bars indicating the 95% confidence interval. Panel A shows per-program slopes with respect to sequencing depth, with a broken x-axis to accommodate the full range of slopes across responses. Panels B-E show the slopes broken down by each genome-architecture covariate: (B) genome size, (C) gene density, (D) GC content, and (E) CDS-per-gene ratio. Colors indicate each program: ABySS short, SPAdes short, Canu, Flye, Hifiasm, Miniasm, NextDenovo, Raven. Shape indicates the significance level: square p ≥ 0.05, triangle p < 0.05, circle p < 0.001.

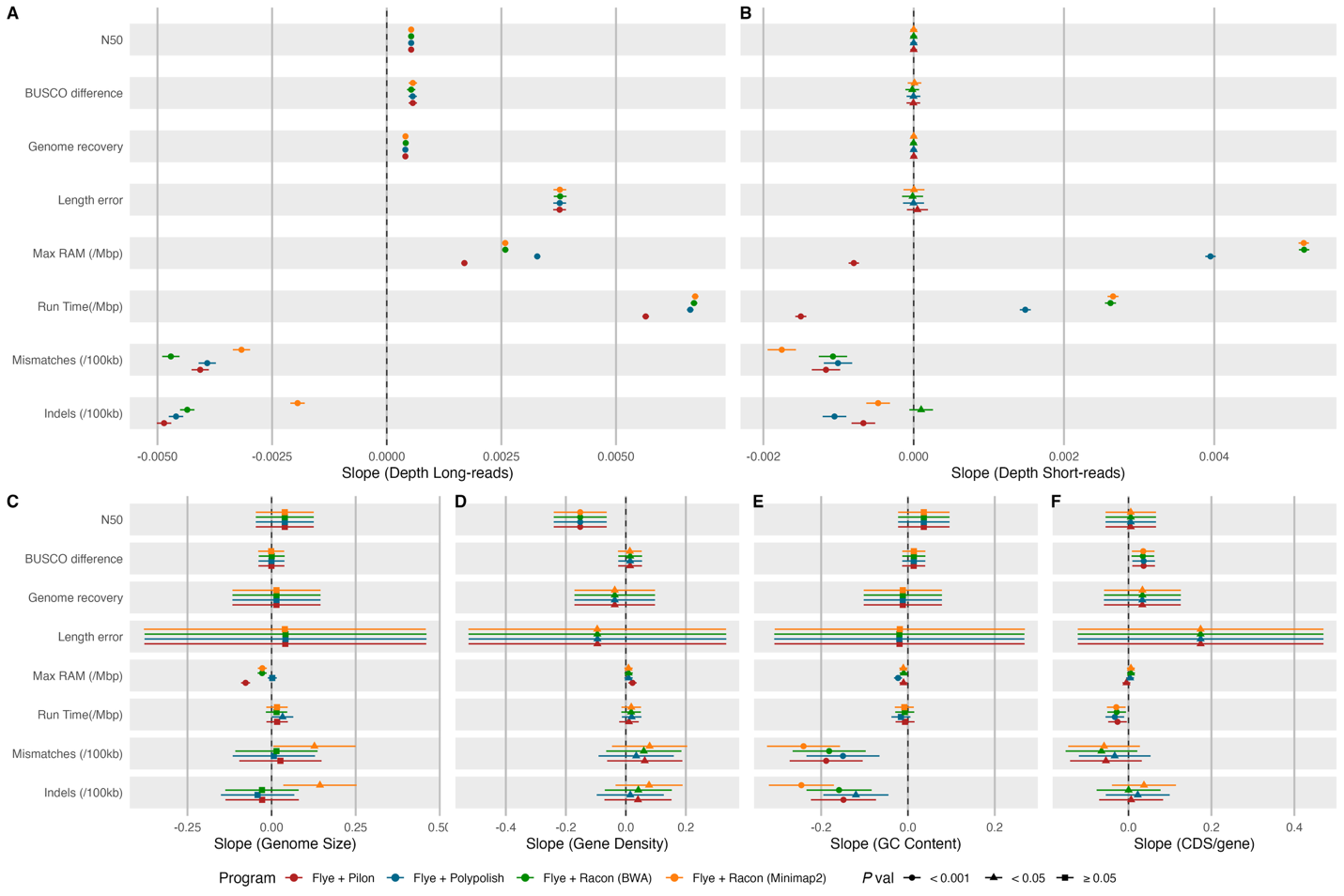

Supplemental Figure 5. Forest plot of per-program slopes for continuous predictors in the simulated polished-strategy model. Slopes were estimated via emmeans::emtrends, with points representing the per-program slope and horizontal bars indicating the 95% confidence interval. Panel A shows per-program slopes with respect to long-reads depth, and Panel B to short-reads, with a broken x-axis to accommodate the full range of slopes across responses. Panels C-F show the slopes broken down by each genome-architecture covariate: (C) genome size, (D) gene density, (E) GC content, and (F) CDS-per-gene ratio. Colors indicate each program or program combination: Flye + Pilon, Flye + Polypolish, Flye + Racon (BWA), Flye + Racon (Minimap2). Shape indicates the significance level: square p ≥ 0.05, triangle p < 0.05, circle p < 0.001.

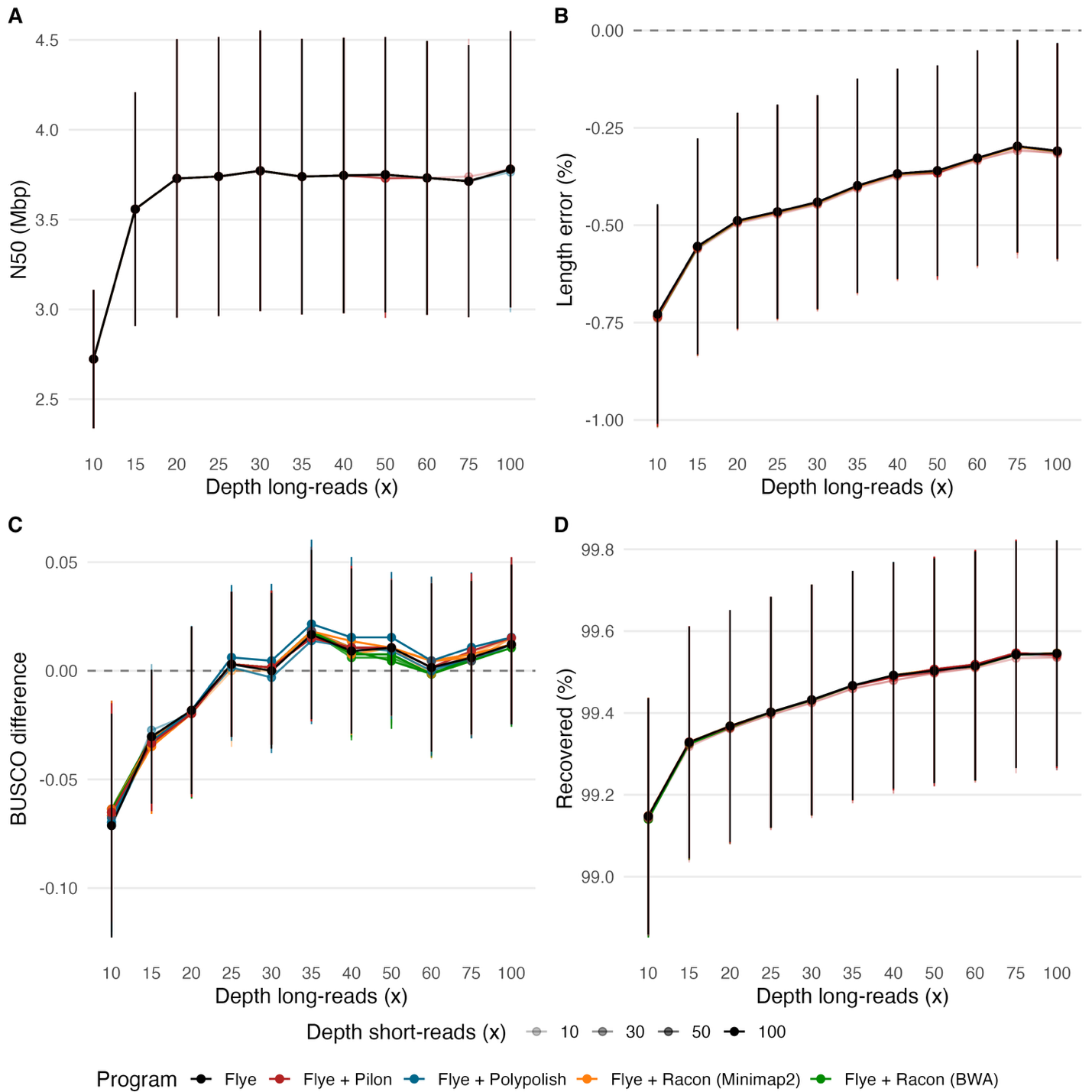

Supplemental Figure 6. Assembly quality metrics for polished Flye assemblies. Panels show (A) N50, (B) length error, (C) BUSCO difference, and (D) recovered percentage for each program. X-axis indicates long-reads depth, and alpha transparency indicates short-reads depth. These results confirm that polishing preserves contiguity and structural characteristics.

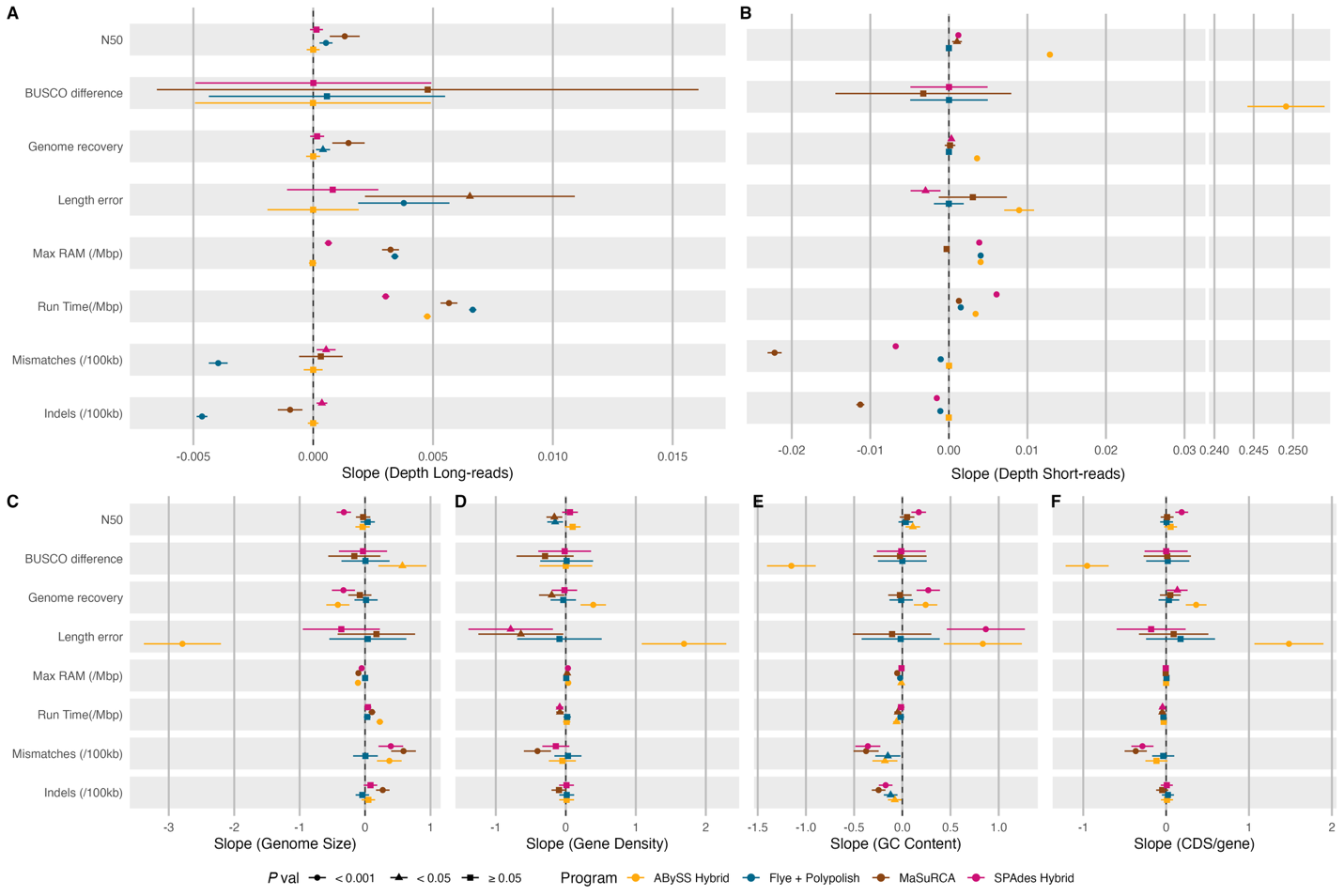

Supplemental Figure 7. Forest plot of per-program slopes for continuous predictors in the simulated hybrid-strategy model. Slopes were estimated via emmeans::emtrends, with points representing the per-program slope and horizontal bars indicating the 95% confidence interval. Panel A shows per-program slopes with respect to long-reads depth, and Panel B to short-reads, with a broken x-axis to accommodate the full range of slopes across responses. Panels C-F show the slopes broken down by each genome-architecture covariate: (C) genome size, (D) gene density, (E) GC content, and (F) CDS-per-gene ratio. Colors indicate each program or program combination: SPAdes hybrid, MaSuRCA, ABySS hybrid, Flye + Polypolish. Shape indicates the significance level: square p ≥ 0.05, triangle p < 0.05, circle p < 0.001.

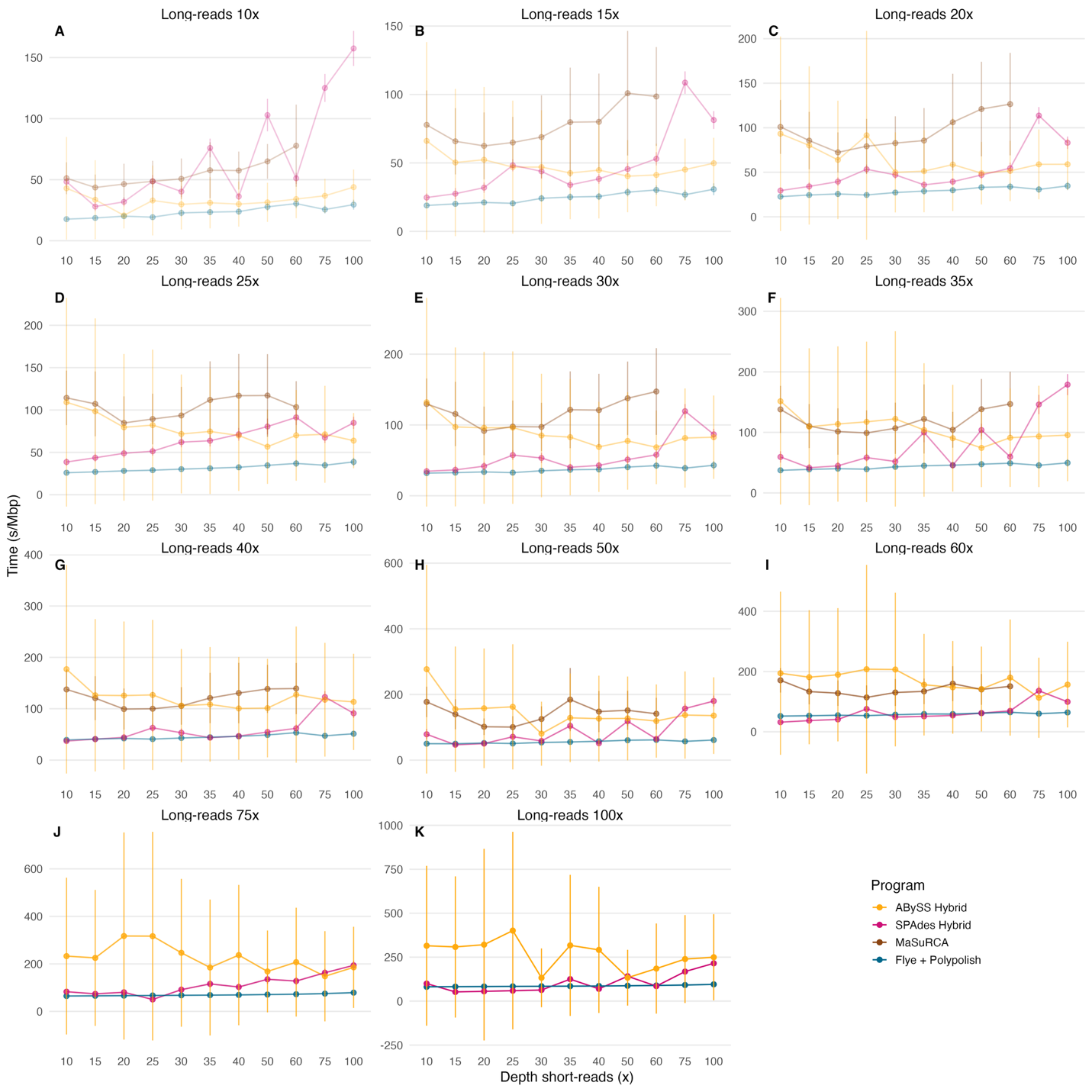

Supplemental Figure 8. Run time per Mbp of assembly for polished and hybrid assembly strategies. Panels indicate distinct long-reads depths: A-10x, B-15x, C-20x, D-25x, E-30x, F-35x, G-40x, H-50x, I-60x, J-75x, K-100x. This is the complete version of Figure 7B. Across depths, runtime increases primarily with long-read coverage and varies substantially by algorithm, with hybrid assemblers showing greater computational escalation than the polished long-read workflow. ABySS Hybrid specially shows grater variation in time at higher depths.

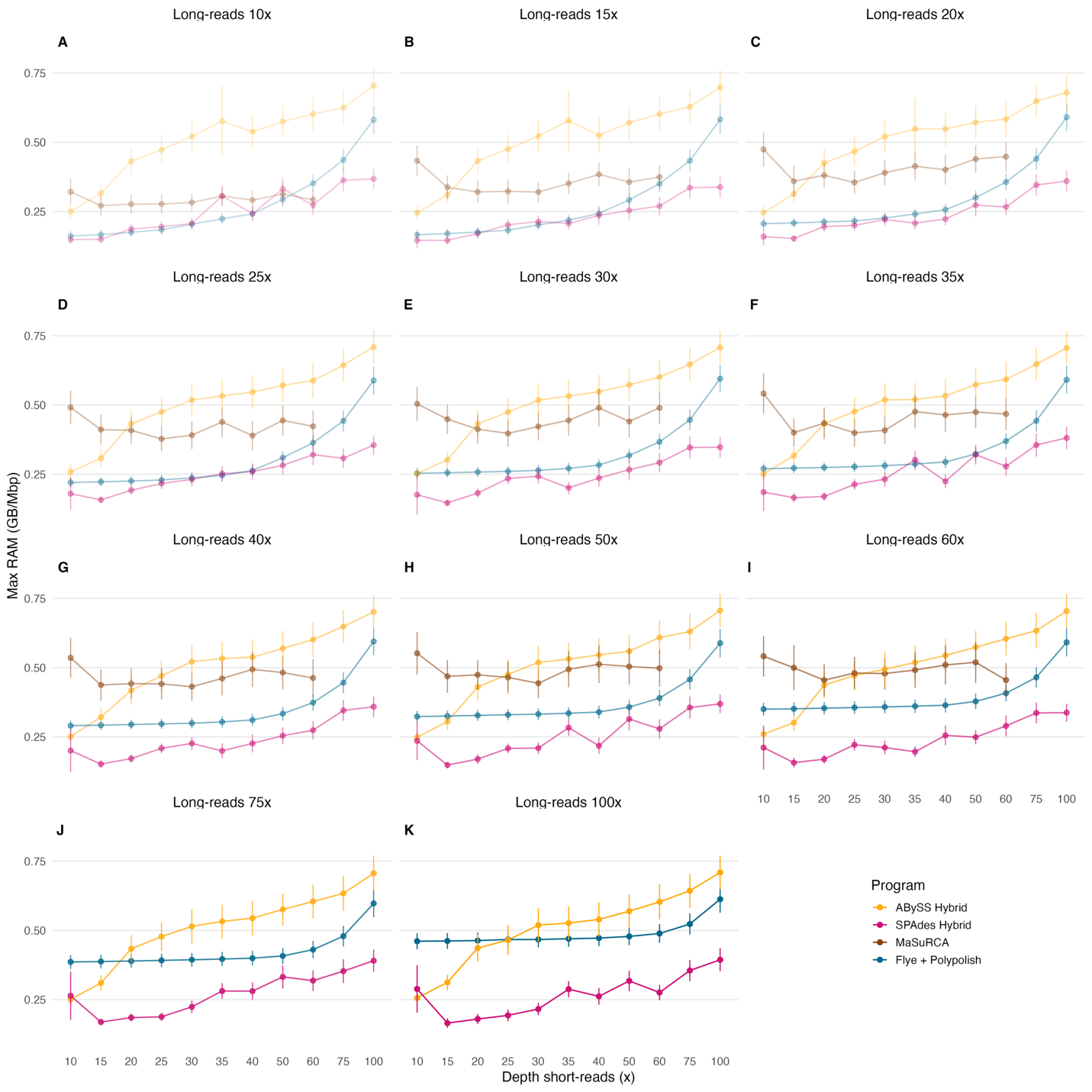

Supplemental Figure 9. Maximum RAM usage per Mbp of assembly for polished and hybrid assembly strategies. Panels indicate distinct long-reads depths: A-10x, B-15x, C-20x, D-25x, E-30x, F-35x, G-40x, H-50x, I-60x, J-75x, K-100x. This is the complete version of Figure 7A. Across coverage levels, memory demand increases primarily with long-read depth and is consistently highest for hybrid assemblers, whereas the polished long-read workflow maintains comparatively stable and lower RAM requirements.

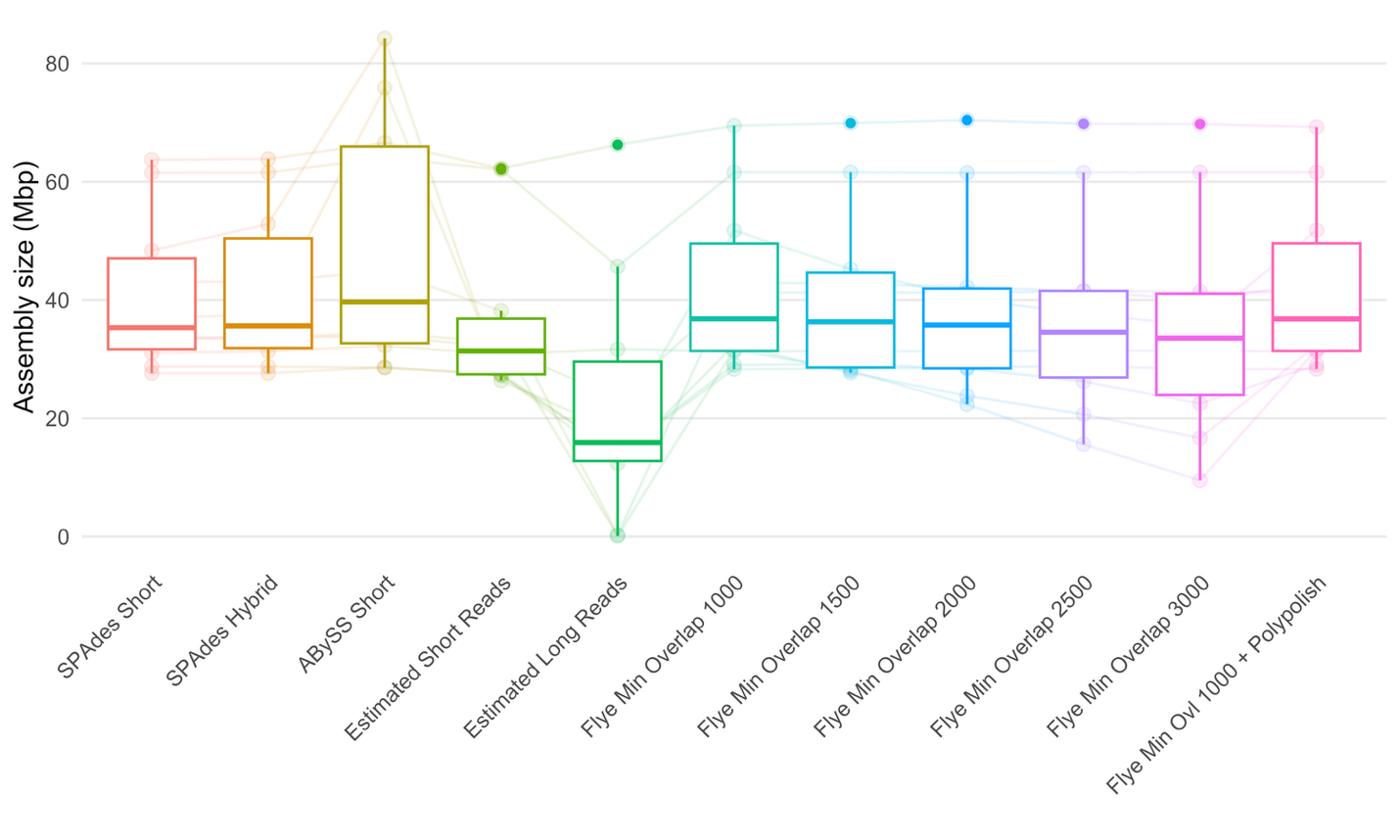

Supplemental Figure 10. Final and estimated assembly size for different programs using the original (not subsampled) long and short reads. Estimation was done using KMC+GenomeScope2 as described in Methods. Short-read–based genome size estimates most closely matched final assemblies, whereas long-read estimates tended to be lower and assembly size varied with assembler strategy and Flye overlap parameters.

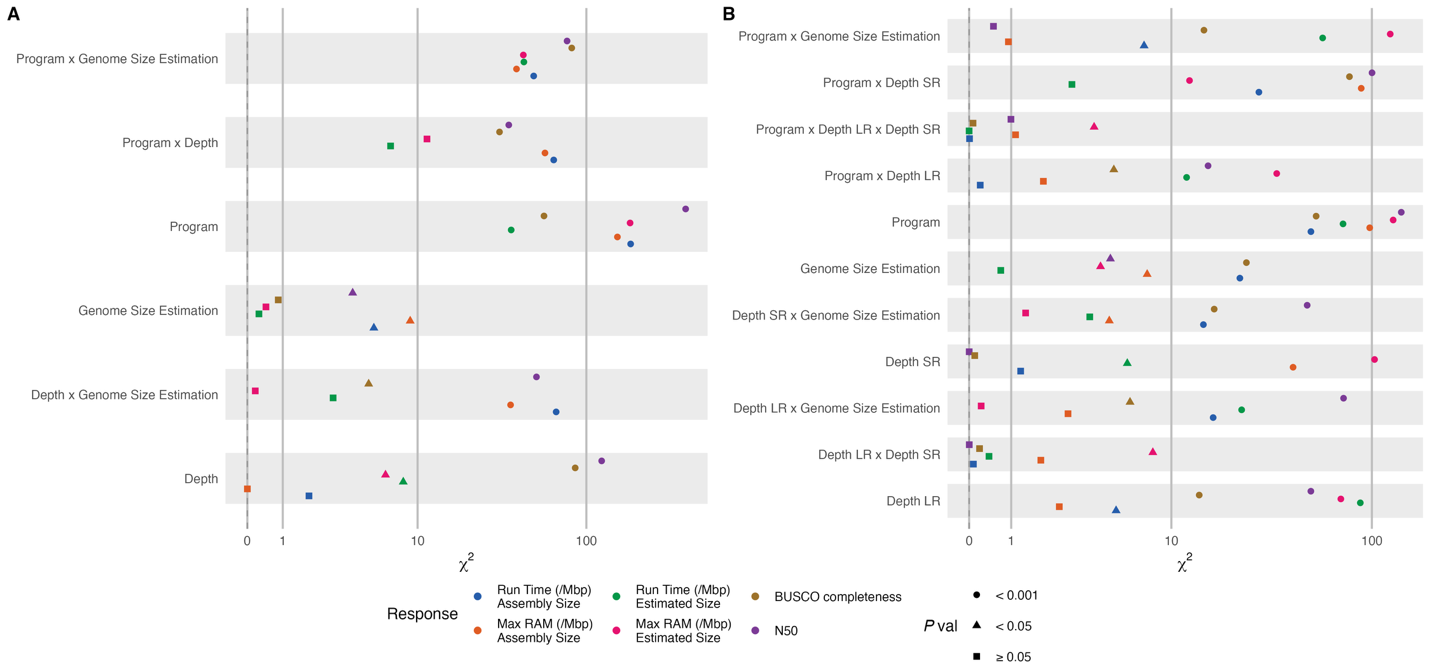

Supplemental Figure 11. Forest plot of Type III Wald χ² statistics for all fixed-effect terms across empirical mixed-effects model. Panels indicate sequence strategy: (A) single sequencing, (B) hybrid sequencing. Color indicates the response and shape indicates the significance level: square p ≥ 0.05, triangle p < 0.05, circle p < 0.001. Points represent the χ² statistic for one fixed-effect term in one response model, plotted on the pseudo-log scaled x-axis.

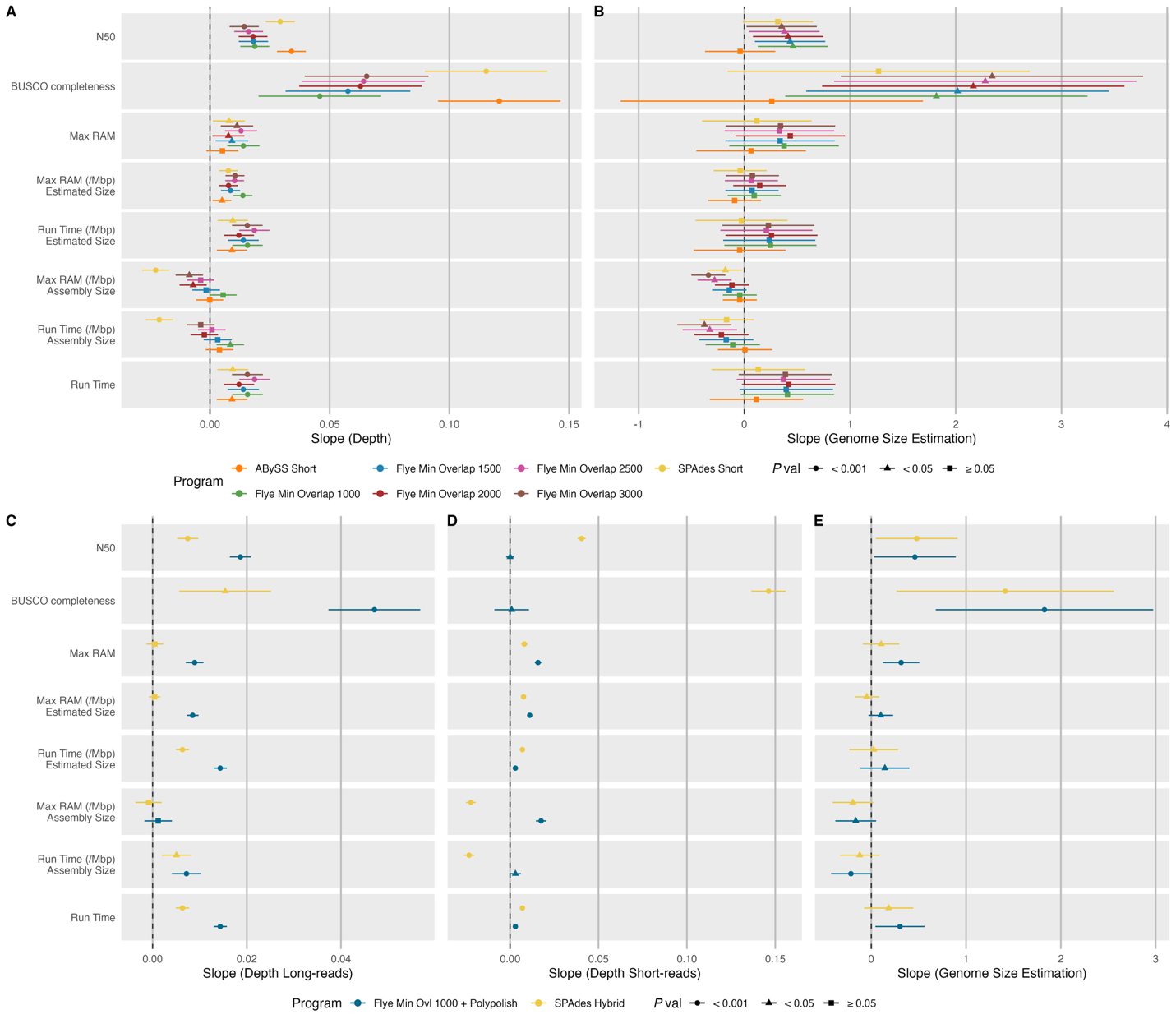

Supplemental Figure 12. Forest plot of per-program slopes for continuous predictors in the empirical dataset. Points represent the per-program slope and horizontal bars indicating the 95% confidence interval. Panels A and B display single-strategy and Panels C-E hybrid strategy. Panels A, C, and D shows per-program slopes with respect to sequencing depth, and Panel B and E shows with respect to the per-isolate genome size estimate. Y-axis corresponds to each response: N50, BUSCO completeness, peak RAM (not normalized), RAM per Mbp of assembly output, RAM per Mbp of proxy genome size, runtime per Mbp of assembly output, runtime per Mbp of proxy genome size, and total elapsed runtime. Colors indicate each program, and shape indicates the significance level: square p ≥ 0.05, triangle p < 0.05, circle p < 0.001.

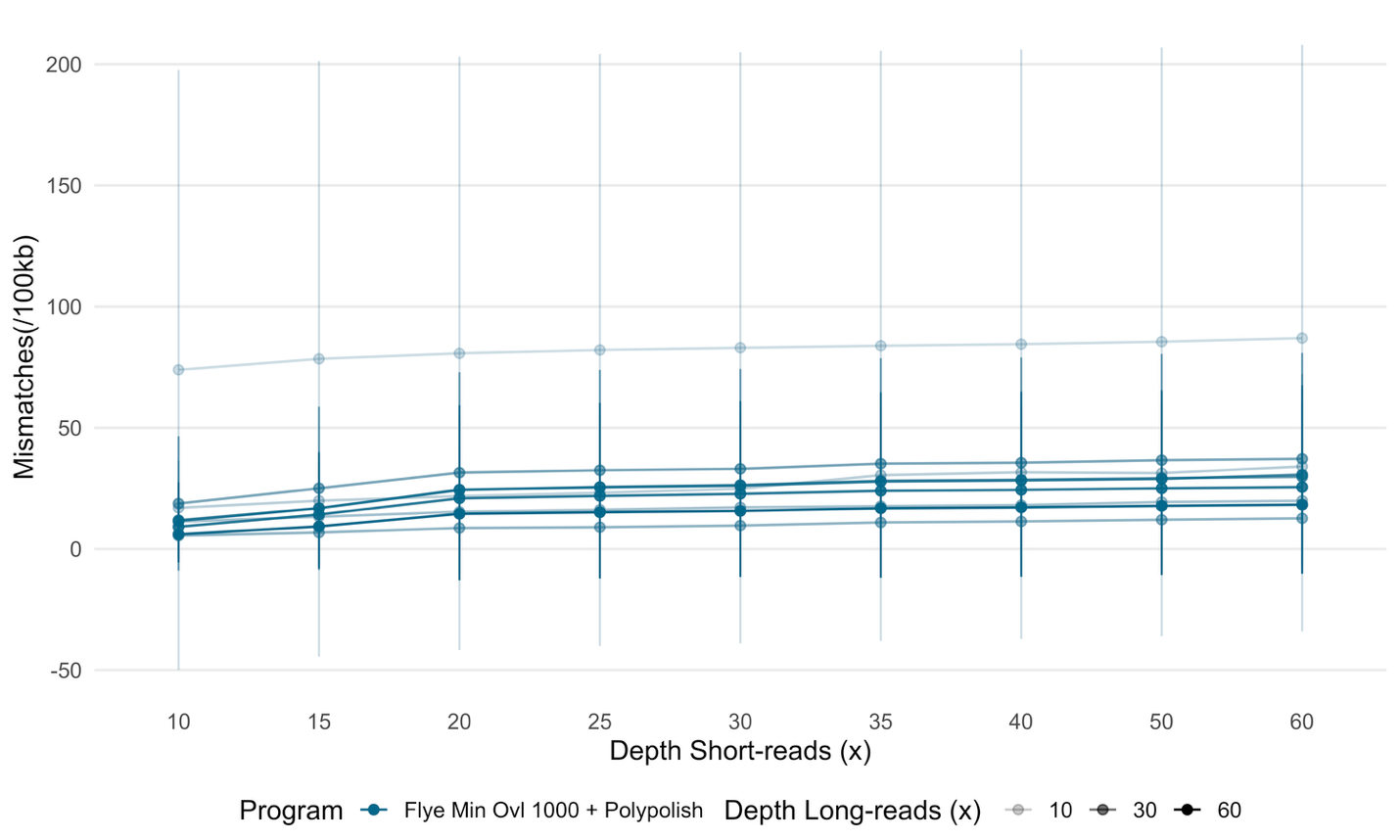

Supplemental Figure 13. Empirical data polished strategy Mismatches per 100Kbp for Flye assemblies with minimum overlap set to 1000 and Polypolish. Lines represent the mean mismatches per 100Kbp for different short-reads depths (x-axis) and long-reads depths (color alpha). Error bars represent 95% confidence intervals around the mean. Mismatch rates decline and stabilize with increasing short-read depth, with most accuracy gains at ~20-30x short-read coverage regardless of long-read depth.
